# Steady-state water exchange in neural tissue is primarily passive and through the phospholipid bilayer

**DOI:** 10.1101/2024.12.12.628254

**Authors:** Nathan H. Williamson, Rea Ravin, Teddy X. Cai, Julian A. Rey, Peter J. Basser

**Author notes:** Conceptualization, NHW, RR, TXC, JAR, and PJB; Methodology, NHW, RR, and TXC; Investigation, NHW and RR.; Writing – Original Draft, NHW; Writing – Review & Editing, NHW, RR, TXC, JAR, and PJB; Funding Acquisition, PJB; Supervision, PJB. The authors have no financial interests to disclose. The views, information or content, and conclusions presented do not necessarily represent the official position or policy of, nor should any official endorsement be inferred on the part of, the Uniformed Services University, the Department of War, the U.S. Government or the Henry M. Jackson Foundation for the Advancement of Military Medicine, Inc. N.H.W. contributed equally to this work with R.R.

## Abstract

Water molecules exchange incessantly across cell membranes and between intracellular compartments, but the dominant steady-state transport pathways, and whether they are active or passive, remain unclear. Low-field, high-gradient diffusion exchange spectroscopy (DEXSY) nuclear magnetic resonance (NMR) measurements on viable *ex vivo* neonatal mouse spinal cords show that water exchange is primarily passive. The apparent exchange rate constant (AXR) depends on osmotic conditions because it reflects multiple exchange pathways, each weighted by the exchanging compartments’ volume fractions. A faster transmembrane path that becomes more visible with increasing extracellular space (ECS) fraction has a high activation energy but is ion-independent, suggesting passive transport through the phospholipid bilayer but not active or passive transport through co-transporter or channel proteins. A slower pathway which dominates when the extracelluar space shrinks has a low activation energy, consistent with geometric exchange between intracellular environments. Moreover, we show how DEXSY can be used to non-invasively measure tonicity in tissue, and inform us about the status of the tissue milieu. These findings may inform future translation to clinical MRI.

**Significance:** We use advanced nuclear magnetic resonance methods to address two unanswered questions in cellular biology: How does water exchange between tissue microenvironments under steady-state conditions, and do these processes involve active water cycling?

---

Water is the solvent of life, yet details of its movement across cell membranes remains an enigma. The dominant transmembrane pathways and whether they involve active or passive transport remain unclear.(1) Noninvasive MRI techniques can image water exchange,(2–5) but unlocking their full diagnostic potential requires a deeper understanding of the mechanisms at play.

Among techniques for probing water exchange, diffusion-based NMR methods are emerging as the most translatable to *in vivo* MRI of the central nervous system (CNS), compared with contrast agent–based approaches.(6) By encoding water displacements with magnetic field gradients, apparent diffusion coefficients (ADC) indirectly reflect cellular microstructure. With two gradient encoding periods separated by a mixing time, diffusion exchange spectroscopy (DEXSY) reveals water movement between compartments with distinct ADC values.(7) With gradients tuned appropriately, intracellular water is more restricted and has a lower ADC than extracellular water,(8) allowing the apparent exchange rate constant (AXR) to register transmembrane exchange,(9) as it is proportional to the membrane surface-to-volume ratio (SVR) and its diffusive permeability to water.(10, 11)

Passive water exchange involves water diffusing through the membrane lipid bilayer or channels, driven only by thermal motion.(12) Sensitivity of exchange to active ion transport inhibitors led to the “active water cycling” (AWC) hypothesis in which water moves in tandem with ions through transporters.(13– 15) Unlike contrast agent-based methods, diffusion-based MR methods can be additionally sensitized to diffusion-mediated geometric exchange between compartments with differing sizes and orientations relative to the gradient direction.(16) While these suggest AXR can be a biomarker for both metabolism and microstructure, the factors governing AXR in geometrically-complex, metabolically-active tissues remain unclear.(17–19)

Active, passive, and geometric water exchange mechanisms potentially are all present in CNS gray matter. Active water cycling (AWC) in gray matter was inferred from contrast-agent studies showing that ouabain inhibition of the Na^+^/K^+^–ATPase strongly reduces water exchange rate constants.(14) This was later supported by low-field, high-gradient DEXSY measurements showing a 70% reduction in AXR.(20) Studies of isolated lipid components suggest that passive membrane permeability could be high(21, 22) and, coupled with the high SVR,(23, 24) could account for fast AXR. Geometric exchange between cell soma and branching processes can also be significant.(24, 25) Simultaneous transmembrane and geometric exchange produce multi-site exchange, introducing cell volume dependencies that simulations showed could account for the 70% AXR reduction.(26) Importantly, active ion transport maintains cell volume,(27) making it difficult to disentangle its alleged direct effect on transmembrane water exchange from an indirect effect on AXR via changes in cell volume. To resolve this, it is necessary to control for tonicity, allowing for perturbations of active ion transport independent of cell volume.

Tonicity is defined by the osmolarity difference of intracellular and extracellular solutes, taking their permeability into account. This inherently affects extracellular space (ECS) and intracellular space (ICS) volume fractions. Intracellular impermeants (metabolites, proteins, nucleic acids, etc.) and cations that accompany them due to their net-negative fixed charges create a persistent osmotic load.(27), which is normally balanced by ions partitioned through active and downstream passive transport processes. In animal cells, the Na^+^/K^+^–ATPase is the primary active transporter. When the Na^+^/K^+^– ATPase is inhibited, the balance is lost and extracellular hypotonicity causes cells to swell. However, in *ex vivo* experiments, osmolytes such as sucrose or mannitol can be added to the bathing media to control tonicity.(23) While tonicity is a critical determinant of water homeostasis, it is difficult or (previously) impossible to measure non-invasively.

In this low-field, high-gradient DEXSY NMR study of viable *ex vivo* spinal cord, we isolate the effects of active ion transport inhibition from those of tonicity changes. Our results indicate that transmembrane water exchange in this system does not involve significant active water cycling (AWC). The data are consistent with passive multi-site exchange, with fast transmembrane exchange between ECS and ICS, and slower geometric exchange between ICS compartments. Under isotonic and hypertonic conditions (i.e., shrunken cells), AXR is fast and exhibits a high activation energy, consistent with passive transmembrane exchange through the phospholipid bilayer. In contrast, hypotonic conditions (i.e., swollen cells) yield slower AXR with a lower activation energy, suggesting water movement through aqueous pathways such as geometric exchange between intracellular regions. The intrinsic permeability of each pathway remains constant, but their relative contributions to the AXR shift with tonicity through changes in ECS volume fraction.

## Results

### Active ion transport inhibition has a large effect on AXR and simultaneously a smaller effect on ADC

The neonatal mouse spinal cord consists mostly of gray matter and little myelinated white matter(28, 29). The microstructure is characterized by the presence of branching neuronal(30) and glial(31) cellular processes with sub-micron radii, and cell bodies (soma) with radii ranging from a few microns to tens of microns.(24) The strong 15.3 T/m static gradient defines the length scale of 800 nm, beyond which diffusing spins dephase and hence restriction by the plasma membranes of cell processes on the the length scale of the cell processes that occu restricted tiffof the measurement measurement sensitivity of what is length scales cause signals to is well-tuned to the length scales of cell processes which occupy the majority of the tissue. In Supporting Information (SI) we show that ADC is linearly correlated to diffusion and DEXSY MR-derived volume fractions (SI 1 and Figs. S2 and S5). This could arise from a simplified two-component model, where the that ADC is inversely proportional to cell volume, which is expected, due to the high gradient strength defining a sensitivity to restriction length scales of roughly 1 micron(32, 33).

AXR and ADC_*y*_ were collected during perturbations with 2, 10, and 100 *µ*M ouabain. Here, the subscript “*y*” refers to the gradient (and hence diffusion encoding) direction, which is perpendicular to the orientation of the spinal cord. 100 *µ*M ouabain is expected to maximally inhibit the Na^+^/K^+^–ATPase, while 2 and 10 *µ*M ouabain are expected to result in partial inhibition.(34) Effects are compared to baseline recordings. Samples maintain stable AXR and ADC_*y*_ for many hours under normal conditions (Figs. 1G, S1D, and Ref. 20) and have been found to be electrically-active afterwards.(32)

**Fig. 1.**
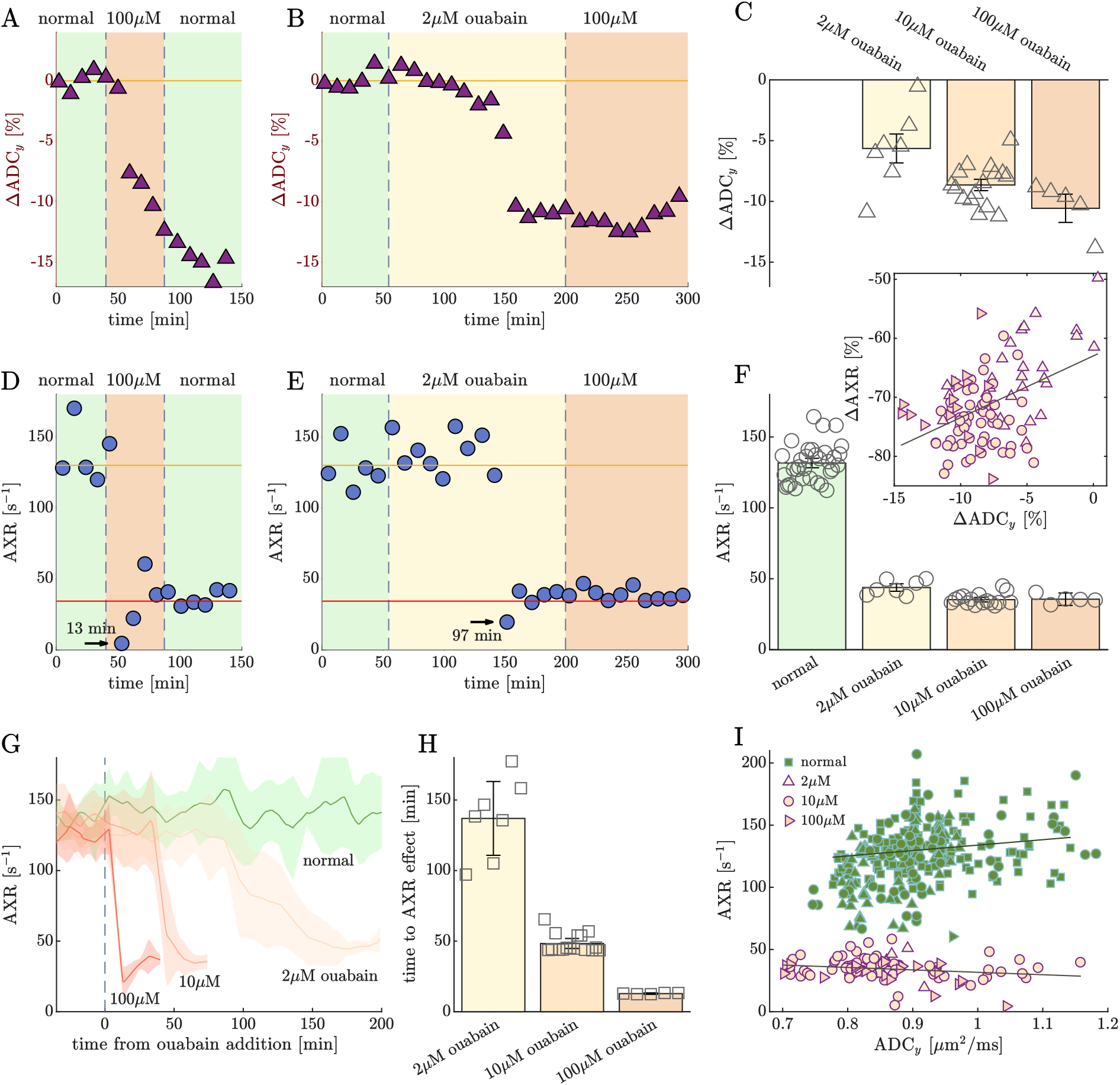
Active ion transport inhibition has a large effect on AXR and simultaneously a smaller effect on ADC. A,B,D,E) Representative recordings on individual samples show how ouabain in (A,D) high (100 *µ*M) and (B,E) low (2 *µ*M) concentrations affects (A,B) ADC in the *y*-direction (presented as % change from baseline, ΔADC_*y*_ = 100 × (ADC_*y*_ − ADC_*y,baseline*_)/ADC_*y,baseline*_) and (D,E) AXR. In (D) and (E), arrows and text show when AXR decreased below 60 s^−1^ and the time from ouabain addition to the effect. C,F) Bar graphs of (C) ΔADC_*y*_ and (F) AXR, showing the mean across all measurements (bar height), 95% CI of the mean (whiskers), and mean values from each sample (open symbols) for normal, 2, 10, and 100 *µ*M ouabain conditions (*n* =33, 7, 16, and 5, respectively). For ouabain conditions, data was compiled from the first 3 to 5 measurements after an effect was observed and did not incorporate data from additional treatments or doses. F-inset) The correlation between ΔADC_*y*_ and Δ*k* for all measurements in the 2, 10, and 100 *µ*M ouabain-treated groups (right-pointing triangles, circles, and up-pointing triangles, respectively). G) Mean (solid lines) and standard deviation (shaded regions) of AXR recordings from a control group under normal conditions (green) and groups treated with 2, 10, and 100 *µ*M ouabain (lightest to darkest colors) plotted against the time after addition of ouabain. H) Bar chart of the time from ouabain addition to the time at which AXR dropped below 60 s^−1^. I) Correlation plot of ADC_*y*_ vs. AXR. Data collected before ouabain treatment makes up the normal group (green squares), and data collected after ouabain treatment was further split into AXR < 60 s^−1^ (pink symbols, see legend) and AXR > 60 s^−1^ (same symbols, but green) groups. See Fig. S2 for additional metrics and correlations.

Ouabain simultaneously decreased ADC_*y*_ and AXR (Fig. 1 and Fig. S1). ADC_*y*_ and AXR were affected precipitously, within 11 minutes after the addition of 100 *µ*M ouabain (Fig. 1A,D,G,H). At 10 *µ*M ouabain, the effect was delayed but still precipitous (Fig. 1G,H). At 2 *µ*M ouabain, the effect was delayed further but occurred on average within 150 min of addition (Fig. 1B,E,G,H). The effect of 2 *µ*M was precipitous in some cases (Fig. 1B,E) but less so overall (Fig. 1G). Subsequent exposure to higher ouabain concentrations did not produce any further effect (Fig. 1B,E). Nevertheless, the initial dose’s impact on ADC_*y*_ was influenced by the concentration (Fig. 1C). 2, 10 and 100 *µ*M ouabain decreased ADC_*y*_ by 5.6 ± 3.1%, 8.6 ± 1.8%, and 10.5 ± 2.2%, respectively (significantly different between groups, *p* < 0.001, see Fig. 1C). In comparison, AXR was much more affected by ouabain, but it depended less on the dosage. AXR is 131 ± 19 s^−1^ under normal conditions. Ouabain reduced AXR by 72 ± 7% (averaged across the three concentration groups, Fig. 1F). 2 *µ*M ouabain reduced AXR to 43.8 ± 6.8 s^−1^ (significantly different from the other concentration groups, *p* < 0.001). 10 and 100 *µ*M ouabain reduced AXR to 35.2 ± 6.0 and 35.6 ± 8.2 s^−1^, respectively (not significantly different, *p* = 0.857). The similar precipitous but delayed effect across dosages (with only a small difference at the lowest 2 *µ*M concentration) is reminiscent of an “all-or-none” effect. The coincident effect on ADC_*y*_ suggests that cells swelled when the drop in AXR occurred. Simulations also predict that deactivating the Na^+^/K^+^–ATPase causes cell swelling and membrane depolarization due to ions repartitioning (Fig. S3).(26)

We tested whether there was a correlation between AXR and ADC_*y*_. First, the relative change from baseline of ΔADC_*y*_ and Δ*k* are plotted for all measurements in the ouabain treated groups (Fig. 1F inset). The correlation is significant but weak (correlation coefficient *cc* = 0.45, *p* < 0.001) and is insignificant when the 2 *µ*M data is omitted (*p* = 0.7). A linear fit of the data yields ΔAXR = 1.05 × ΔADC_*y*_ − 63.0 % with a *y*-intercept shifted well below 0, suggesting a nonlinear relationship which will be elucidated below. Next, the data is grouped based on whether ouabain was administered and correlations between the absolute values of ADC_*y*_ and AXR are presented. The two groups do not share the same correlation (Fig. 1I). ADC_*y*_ and AXR showed a weak positive correlation under normal conditions (*cc* = 0.20, *p* = 0.01) and a weak negative correlation after ouabain took effect (*cc* = − 0.21, *p* = 0.03). Note that the correlation seen when plotting ADC_*y*_ as a percent change from baseline appears much different than when plotting the absolute value (compare Fig. 1F inset to Fig. 1I). This is due to differences between samples which are accounted for when normalizing to the baseline (compare Fig. 1C to Fig. S1C).

These results are consistent with previous studies,(14, 20) and could be construed as evidence for AWC. However, it is difficult to completely rule out the possibility that these results could be explained by more complicated microstructural effects since active ion transport and cell volume are normally interlinked and are both affected by Na^+^/K^+^–ATPase inhibition.

### AXR is not directly linked to ion transport, voltage, or active water cycling

Certain cotransporters have been shown to co-transport water with ions.(35) Similarly, the AWC mechanism, which involves water actively cycling along with ions through the Na^+^/K^+^–ATPase and other downstream transporters, was proposed to explain the effect of ouabain on AXR.(14, 15) Here we test this AWC hypothesis by studying the effects of Na^+^/K^+^–ATPase inhibition independent from volume changes. Since the Na^+^/K^+^– ATPase pumps ions to control tonicity and cell volume under normal conditions,(27) we used an osmolyte to compensate for tonicity changes induced by Na^+^/K^+^– ATPase inhibition. Specifically, we used an aCSF media in which the major ionic component, 128.35 mM NaCl, was replaced with 256.7 mM sucrose (Fig. 2A,B). This was inspired by electrophysiology studies that use 0 NaCl, high sucrose media to maintain viability of adult animal brain slices.(36) Compared to normal aCSF, the 0 NaCl, 256.7 mM sucrose media has an equal osmolarity but a higher tonicity because membranes are impermeable to sucrose but are semipermeable to Na^+^ and Cl^−^ (Fig. S3). Note that the media still has 21.58 mM Na^+^ and 7 mM Cl^−^ from salts other than NaCl.

**Fig. 2.**
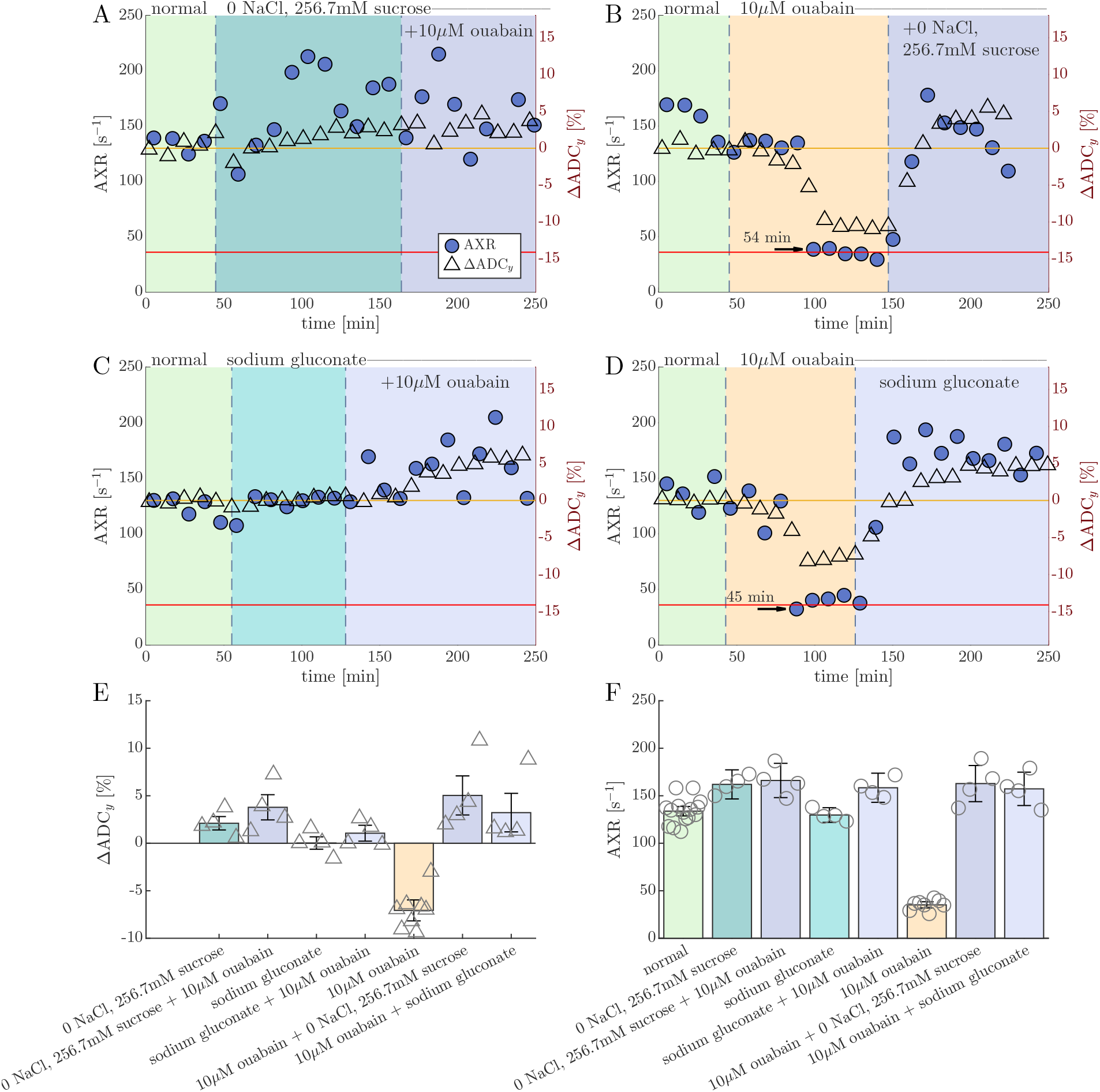
AXR is not directly linked to ion transport, voltage, or active water cycling. A–D) Representative recording of AXR (blue circles) and % change in ADC_*y*_ (open triangles) during experiments involving (A,C) washing the sample from normal media to media in which NaCl was replaced with equiosmolar concentration of (A) sucrose (256.7 mM) or (C) sodium gluconate (128 mM), and then adding 10 *µ*M ouabain, and (B,D) adding 10 *µ*M ouabain on top of normal media, waiting for the effect, and then washing to (B) 0 NaCl, 256.7 mM sucrose media or (D) sodium gluconate media in the presence of 10 *µ*M ouabain. E,F) Box plots of (E) ΔADC_*y*_ and (F) AXR showing the statistics across *n* = 4 samples.

The 0 NaCl, 256.7 mM sucrose media caused ΔADC_*y*_ to increase significantly (*p* < 0.001), by 2.1 ± 1.3%, because its higher tonicity shrunk cells slightly (Fig. 2A,E). If water actively cycles along with ions, then the transmembrane exchange rate constant would decrease when washing to 0 NaCl, 256.7 mM sucrose media because Na^+^/K^+^–ATPase activity is reduced and any potential AWC membrane influx proteins acting as Na^+^ and Cl^−^ cotransporters should be reduced considerably. Instead, AXR increased significantly (*p* < 0.001), by 27 ± 24%, and 10 *µ*M ouabain had no additional effect (*p* = 0.7) (Fig. 2A,F). When ouabain was added first and was present through the whole experiment, the 0 NaCl, 256.7 mM sucrose media subsequently raised AXR significantly above normal levels (*p* < 0.001) (Fig. 2B,F). Clearly, since the effect of inhibiting the Na^+^/K^+^–ATPase can be undone by an osmolyte, AXR is not directly linked to Na^+^/K^+^–ATPase activity. Instead, it is linked to the tonicity, either established by the Na^+^/K^+^–ATPase or external osmolytes.

To distinguish effects related to cell transmembrane voltage from effects related to cell volume, normal aCSF media was washed to media in which the 128.35 mM NaCl was replaced with 128.35 mM sodium gluconate. Gluconate is a monovalent anion and does not pass through the plasma membrane.(37) 128.35 mM sodium gluconate is expected to have the same effect on tonicity as 256.7 mM sucrose. However, the Na^+^/K^+^–ATPase can still be active and voltage can still be maintained in the sodium gluconate media since Na^+^ is still present in high concentration. Simulations show that when the Na^+^/K^+^– ATPase is inactivated, in normal aCSF cells swell and depolarize whereas in 128.35 mM sodium gluconate media cells depolarize without changing volume (Fig. S3).(26) This is because gluconate does not flow into cells following loss of membrane potential as chloride would.

We found that washing to sodium gluconate media had no significant effect on ADC_y_ or AXR (*p* = 0.9 and *p* = 0.6, respectively) (Fig. 2C,E,F). Adding 10 *µ*M ouabain on top of sodium gluconate media caused ADC_y_ to increase significantly (*p* = 0.02), indicating cellular shrinkage. AXR also increased significantly (*p* < 0.001), to values similar to those obtained under 0 NaCl, 256.7 mM sucrose media conditions (*p* = 0.001). The lack of effect when washing to sodium gluconate followed by the effect when adding ouabain could be evidence of cell volume regulation.(38) The data variability increased after ouabain addition, which could be due to slow changes occurring during the measurement. Experiments were repeated with ouabain being added before washing to sodium gluconate media and yielded consistent results (Fig. 2D,E,F).

We conclude from the results above that steady-state transmembrane exchange in this system does not have a significant active component on the 1–300 ms timescales probed by low-field, high gradient DEXSY.

### AXR depends on tonicity

Next, we study the effects of osmolytes (sucrose and mannitol) on ADC_*y*_ and AXR under normal conditions as well as after ouabain administration. This added pressure shrinks cells, increasing the concentration of trapped intracellular impermeants.(27) In viable tissue this also activates regulatory volume increase (RVI) mechanisms, which involve Na^+^ and K^+^ uptake coupled with electroneutral transport of Cl^−^, in-part by reducing Na^+^/K^+^–ATPase activity.(38, 39) These two effects contribute to balancing the osmotic pressure and determining new steady-state cell volumes.

First, the effect of +100 mOsm sucrose was observed on the same sample before and after adding 10 *µ*M ouabain (Fig. 3). ADC_*y*_ followed trends expected for cellular shrinking and swelling; it increased in the +100 mOsm hypertonic conditions, recovered in normal isotonic media, and decreased when ouabain took effect (Fig. 3A,B). In the context of AWC, increased osmolarity would be expected to reduce Na^+^/K^+^–ATPase activity(39) and in this way reduce AXR. In contrast, starting from the normal condition, +100 mOsm increased AXR slightly but significantly (*p* = 0.03), by 14 ± 25%. (Fig. 3C,D). After ouabain took effect, +100 mOsm increased AXR much more, by 233 ± 68%. This further suggests AXR is related to the tonicity. AXR dropped back down when the osmolyte was washed away in the presence of ouabain (Fig. 3C,D).

**Fig. 3.**
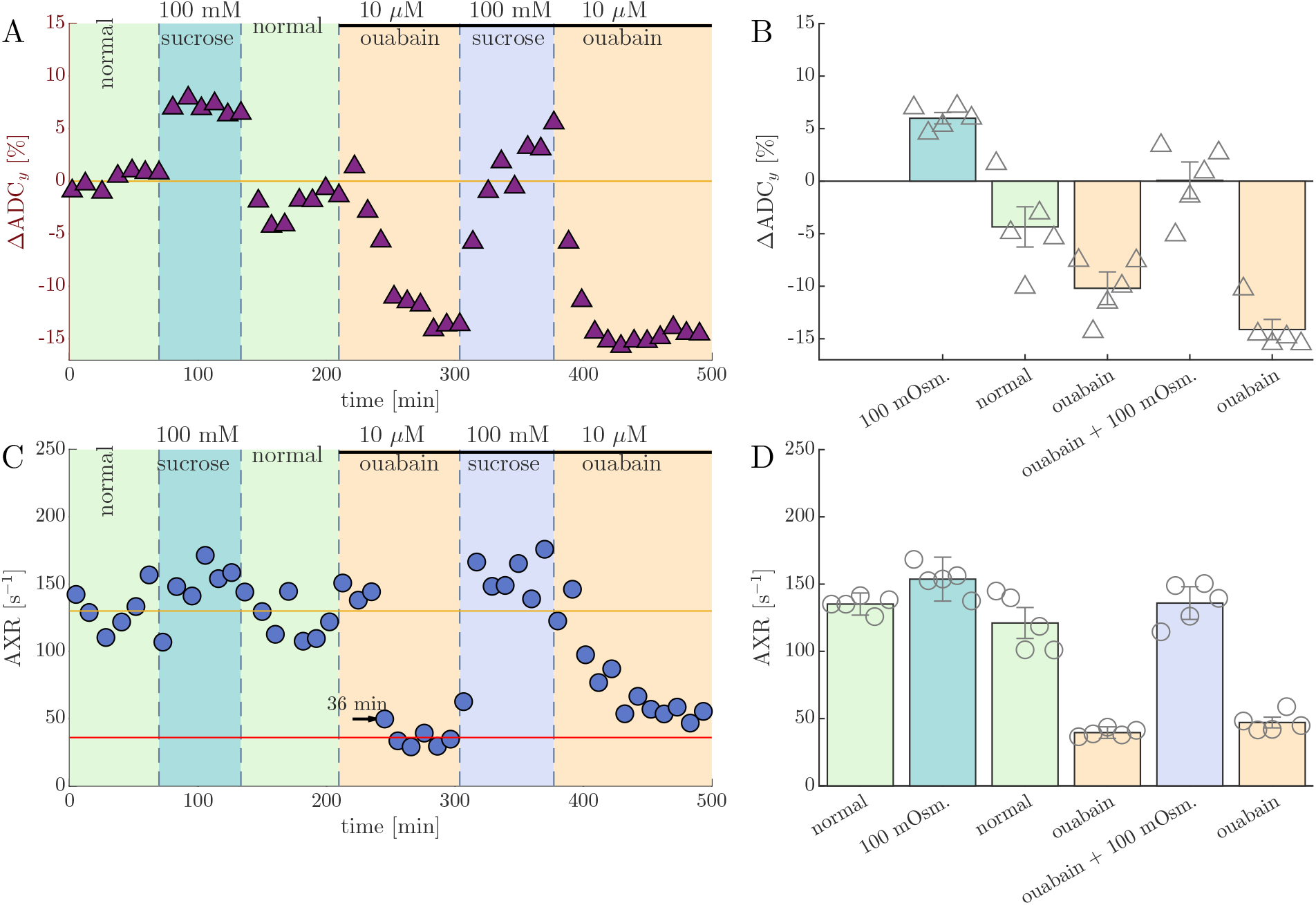
Osmolytes recover AXR. A,C) Representative recordings of (A) % change in ADC_y_ and (C) AXR during perturbations with 100 mM sucrose (an osmolyte) before and after the addition of 10 *µ*M ouabain. The presence of 10 *µ*M ouabain is shown by the solid-black line. B,D) Bar graphs showing the statistics across *n* = 5 samples.

Next, the effects of sucrose dosages in the range of +10 to +200 mOsm were studied on normal samples and on separate samples treated with 10 *µ*M ouabain (Fig. 4A,B). On normal samples, ΔADC_*y*_ and AXR increased only slightly; the effect was insignificant for dosages up to +100 mOsm but became significant at +200 mOsm (Fig. 4A,C,E). For ouabain-treated samples, ΔADC_*y*_ and AXR increased significantly for doses between +30 mOsm and +100 mOsm. The trend changed at +200 mOsm, with ΔADC_*y*_ and AXR decreasing slightly, although not significantly (Fig. 4B,C,E). Comparing ΔADC_*y*_ and AXR between the normal and ouabain-treated groups, both values were significantly lower for ouabain-treated samples at +0 mOsm and similar between groups at +100 mOsm. At +200 mOsm, AXR of ouabain-treated samples was significantly less than that of normal samples (Fig. 4E) but ΔADC_*y*_ remained similar between sample groups (Fig. 4C). The effects of mannitol were also studied on ouabain-treated samples (Fig. 4D). Mannitol (MW= 182 g/mol) is a smaller molecule than sucrose (MW= 342 g/mol) and is also used as a cellular osmolyte.(23) Mannitol and sucrose increased ΔADC_*y*_ to similar values at each concentration (Fig. 4C). This is in spite of sucrose reducing the ADC of pure aCSF media to a greater degree than mannitol due to its greater effect on fluid viscosity (Fig. S4). Mannitol and sucrose also increased AXR similarly across concentrations (Fig. 4E).

**Fig. 4.**
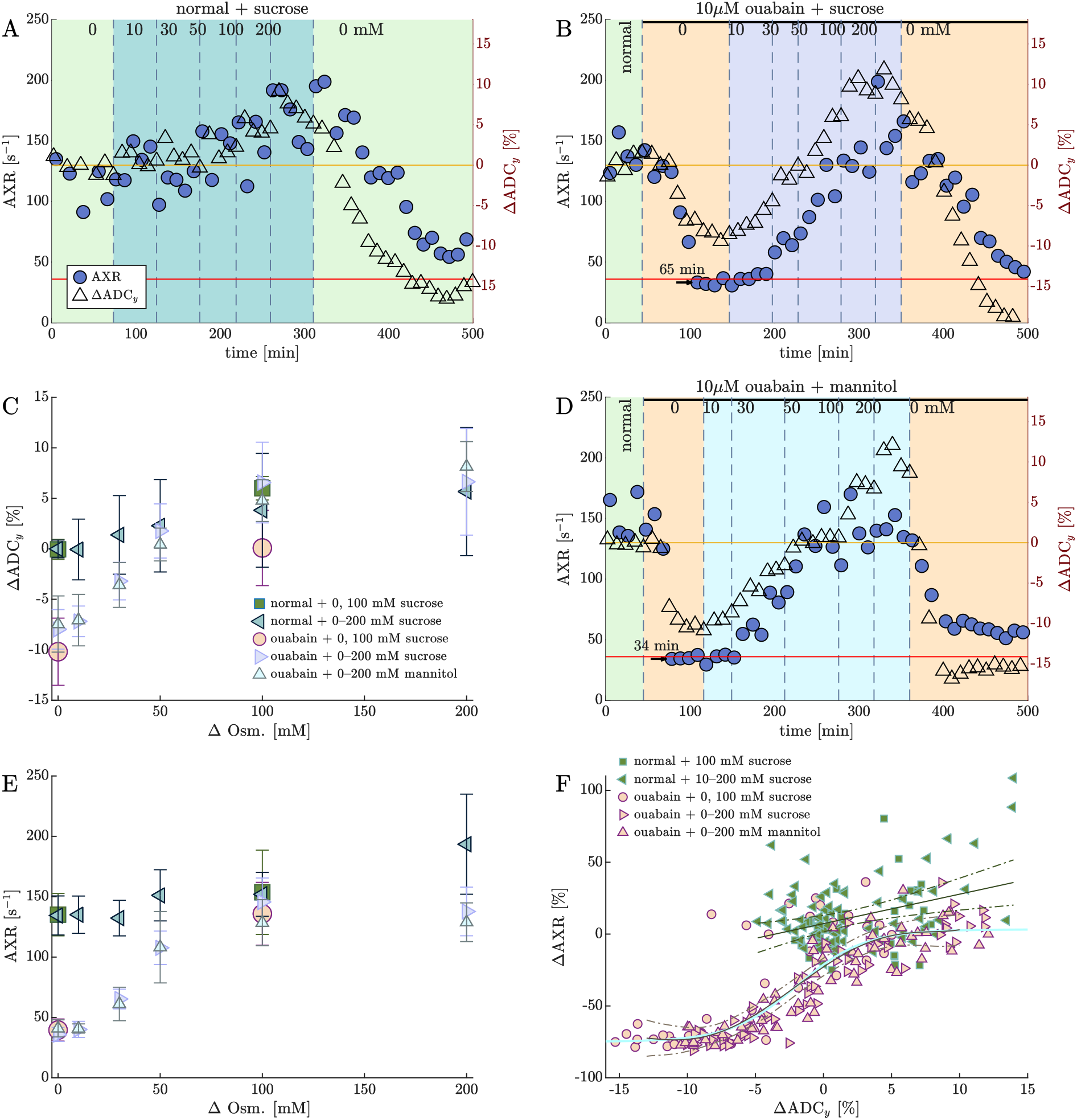
AXR depends on tonicity. A,B,D) Recordings of AXR (left *y*-axis) and ΔADC_*y*_ (right *y*-axis) during osmotic perturbations with (A,B) sucrose or (D) mannitol concentrations 0, 10, 30, 50, 100, 200 mM and back to 0 mM on representative samples (A) starting from normal and (B,D) in the presence of 10 *µ*M ouabain (shown by solid-back line). C,E) The effect of osmolarity on the mean (symbols) and standard deviations (error bars) of (C) ADC_*y*_ and (E) AXR when adding sucrose to samples under normal conditions (*n* = 3), and when adding sucrose (*n* = 3) or mannitol (*n* = 3) to samples treated with 10 *µ*M ouabain (see legend in C). Data from Fig. 3 involving the addition of 100 mM sucrose in a single step pre-ouabain treatment and post-ouabain treatment are also shown. F) The correlation between ΔADC_*y*_ and Δ*k* for all measurements performed on all samples in each group (see legend). Predicted means (solid lines) and 99% CI (broken lines) were found by regression of 1^st^ order (combined normal samples) and 5^th^ order (combined ouabain-treated samples) polynomials, with the latter being compared to a sigmoidal logistics function (aqua blue line). Additional metrics and correlations are presented in Fig. S5.

While effects of 100 mOsm achieved in one large dose vs. incremental doses were consistent (Fig. 4C,E), differences were observed on normal samples during the wash to normal media (comparing Fig. 4A to Fig. 3A,C) and are discussed in SI 2.

We tested whether the effects of osmolytes were similar or different between normal and ouabain-treated groups by examining correlations between ΔADC_*y*_ and ΔAXR (Fig. 4F). The two groups have different correlations and occupy distinct regions of the plot (see distinct model fits and 95% CI). Data from normal samples appears clustered due to osmolytes having little effect. For ouabain-treated samples, ΔADC_*y*_ and ΔAXR show a sigmoidal relationship, being highly correlated at intermediate ΔADC_*y*_ and weakly correlated at low and high ΔADC_*y*_. This behavior is not due to measurement insensitivity because we report lower AXR’s for ouabain-treated samples at 11^°^C (Fig. 5E) and we previously reported higher AXR’s for normal samples at 35^°^C (in Fig. 1B of Ref. 20).

**Fig. 5.**
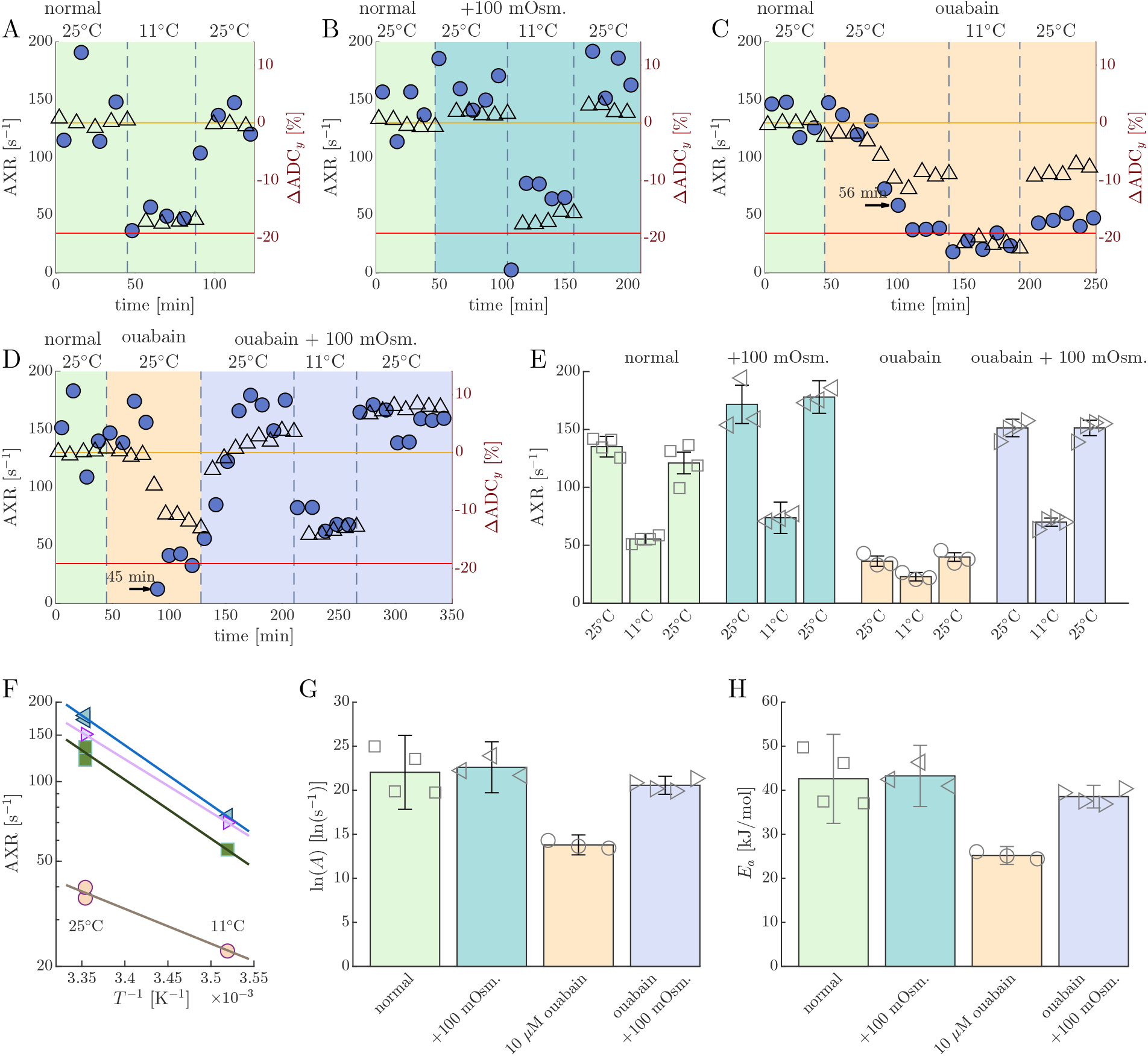
Tonicity affects activation energies of AXR. A–D) Recordings of AXR (blue circles) and ΔADC_*y*_ (open triangles) during temperature perturbation from 25^°^C to 11^°^C back to 25^°^C under (A) normal, (B) +100 mM sucrose, (C) 10 *µ*M ouabain, (D) and 10 *µ*M ouabain +100 mM sucrose conditions. E) Results of AXR recorded at 25^°^C, 11^°^C, and again at 25^°^C on samples under normal (*n* = 4), +100 mOsm (*n* = 3), 10 *µ*M ouabain (*n* = 3), and 10 *µ*M ouabain +100 mOsm (*n* = 4) conditions. F) Arrhenius plot of the inverse of the absolute temperature *T* ^−1^ vs. the average AXR for each condition. Solid lines show average fits of the Arrhenius model (Eq. 1). The slope of the line on the semi-log plot is proportional to *E*_*a*_. (Refer to error bars in E, G, and H for standard deviations of AXR and Arrhenius model parameters.) G,H) Bar graph of (G) the natural log of pre-exponential factor *A* and (H) activation energy *E*_*a*_ estimated from Arrhenius model fits.

While osmolytes restore AXR of ouabain-treated samples to baseline levels, it is unclear whether the same exchange pathways are recovering or if SVR changes are responsible. Measuring activation energies from the temperature dependence of AXR can help distinguish these causes, as they reflect the energy barriers of underlying water exchange pathways, they do not depend on SVR differences between sample groups, and they can be compared across studies.(12)

### Tonicity affects activation energies of AXR but not of ADC

Here, we study samples under normal conditions, treated with 10 *µ*M ouabain, treated with 100 mM sucrose, and treated with 10 *µ*M ouabain + 100 mM sucrose. For each condition, AXR and ADC_*y*_ was recorded at 25^°^C and 11^°^C, and re-recorded at 25^°^C (Fig. 5A–E). AXR recovered when re-recorded at 25^°^C (one-way ANOVA, *p >* 0.05). The dependence of AXR on the absolute temperature *T* is modeled with the Arrhenius equation

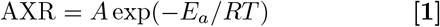

where *A* is the pre-exponential factor, *E*_*a*_ is the activation energy, and *R* is the ideal gas constant (Fig. 5F). We previously studied exchange at three temperatures in spinal cord tissue and did not see deviations from Arrhenius behavior,(20) thus justifying the use of only two temperatures here. Here we find *E*_*a*_ values for AXR are similar between the normal samples (*E*_*a*_ = 42.6 ± 6.4 kJ/mol) and normal +100 mOsm samples (43.2 ± 2.8 kJ/mol), but different from the ouabain-treated samples (25.2 ± 0.8 kJ/mol, Fig. 5H). *E*_*a*_ recovers to 38.5 ± 1.6 kJ/mol when +100 mOsm is added to ouabain-treated samples, indicating that the recovery is related to tonicity but not directly related to Na^+^/K^+^–ATPase activity. The difference in *E*_*a*_ between ouabain-treated samples and the other samples also indicates that SVR changes alone cannot account for the observed AXR differences. Faster AXR is associated with higher *E*_*a*_ (compare panels E and H of Fig. 5). This is unexpected. In a simple two-compartment model for transmembrane exchange, one would expect increased AXR from increased permeability to be associated with *lower E*_*a*_. Later, we show this can arise in multi-site exchange involving exchange pathways that have different activation energies.

Temperature affects water’s thermal motion and self-diffusion coefficient.(40) In tissue, as temperature increases, the diffusion length scale and number of interactions with membrane surfaces increase. The ADC still goes up with temperature, but not as much as it would if diffusion were unimpeded, leading to the *E*_*a*_ being artificially reduced from that of pure water or an aqueous solution.(20) Indeed, values were lower than the value found for water self-diffusion in aCSF media (*E*_*a*_ = 18 kJ/mol).(20) In contrast to AXR, the Arrhenius analysis of ADC_*y*_ yielded similar *E*_*a*_ values between conditions (Fig. 6). This is consistent with the variability in ADC_*y*_ between conditions being simply related to microstructural effects like cellular swelling and shrinking and not to different diffusion mechanisms.

**Fig. 6.**
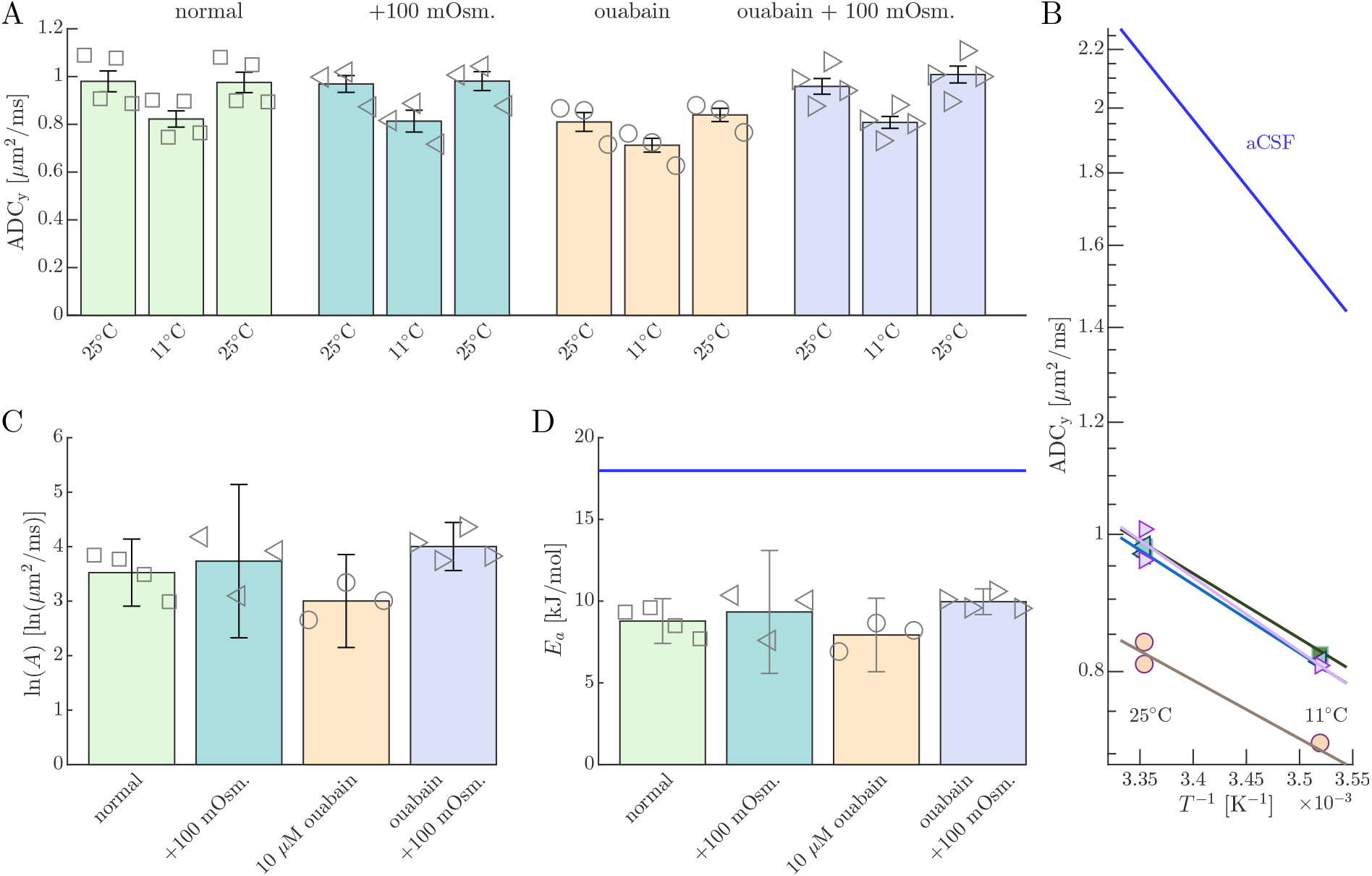
Tonicity does not affect activation energies of ADC. A) Results of ADC_y_ recorded at 25^°^C and 11^°^C and re-recorded at 25^°^C on samples under normal conditions (*n* = 4), after addition of 100 mM sucrose (*n* = 3), after the effect of 10 *µ*M ouabain (*n* = 3), and after the combined effect of 10 *µ*M ouabain and 100 mM sucrose (*n* = 3). B) Arrhenius plot of the inverse of the absolute temperature *T* ^−1^ vs. the average ADC_y_ for each condition and for aCSF (reproduced from Ref. 20). Solid lines show average fits of the Arrhenius model (similar form to Eq. 1). C) Bar graph of the natural log of pre-exponential factor estimates. D) Bar graph of activation energy estimates, compared to the value for aCSF (blue line).

### AXR increasing with activation energy follows enthalpy–entropy compensation

As stated earlier, from a perspective of passive exchange between two sites, it is difficult to explain why faster AXR is associated with higher AXR *E*_*a*_. For instance, if AXR increased due to an increase in membrane permeability, we would expect *E*_*a*_ to decrease. While SVR differences can affect AXR,(11) they should not affect *E*_*a*_.

In general kinetic phenomena, this behavior can arise when the overall rate constant depends on multiple competing pathways with different activation energies (41, 42) and when the relative contributions of these pathways can be modulated by changes in solution composition. (43) This behavior leads to a common linear dependence between *E*_*a*_ and the natural log of *A* across compositions,

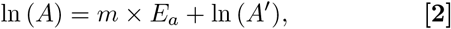

where *m* and *A*′ are constants.(44) This implies that the system under any condition can be described by a modified Arrhenius model of the form

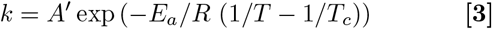

with a crossover temperature *T*_*c*_ = 1/(*m* × *R*) at which AXR values are equal to *A*′. The crossover temperature is a common feature in a more general phenomena termed enthalpy–entropy compensation (EEC) because the concentration-dependent changes in enthalpies and entropies of activation (related to *E*_*a*_ and *A*, respectively) compensate each other (discussed in SI section 3).(41, 43, 45) Consistent with EEC, the *E*_*a*_ and ln(*A*) values were found to be linearly related through Eq. 2 across all conditions (Fig. 7A). This led to estimates of *T*_*c*_ = −17^°^C and *A*′ = 9.5 s^−1^ which, when put into Eq. 3, describes the temperature dependence of AXR for all systems (Fig. 7B). Appearance of EEC suggests a common mechanism exists under all conditions, where *E*_*a*_ and *A* are a function of tonicity alone.(41)

**Fig. 7.**
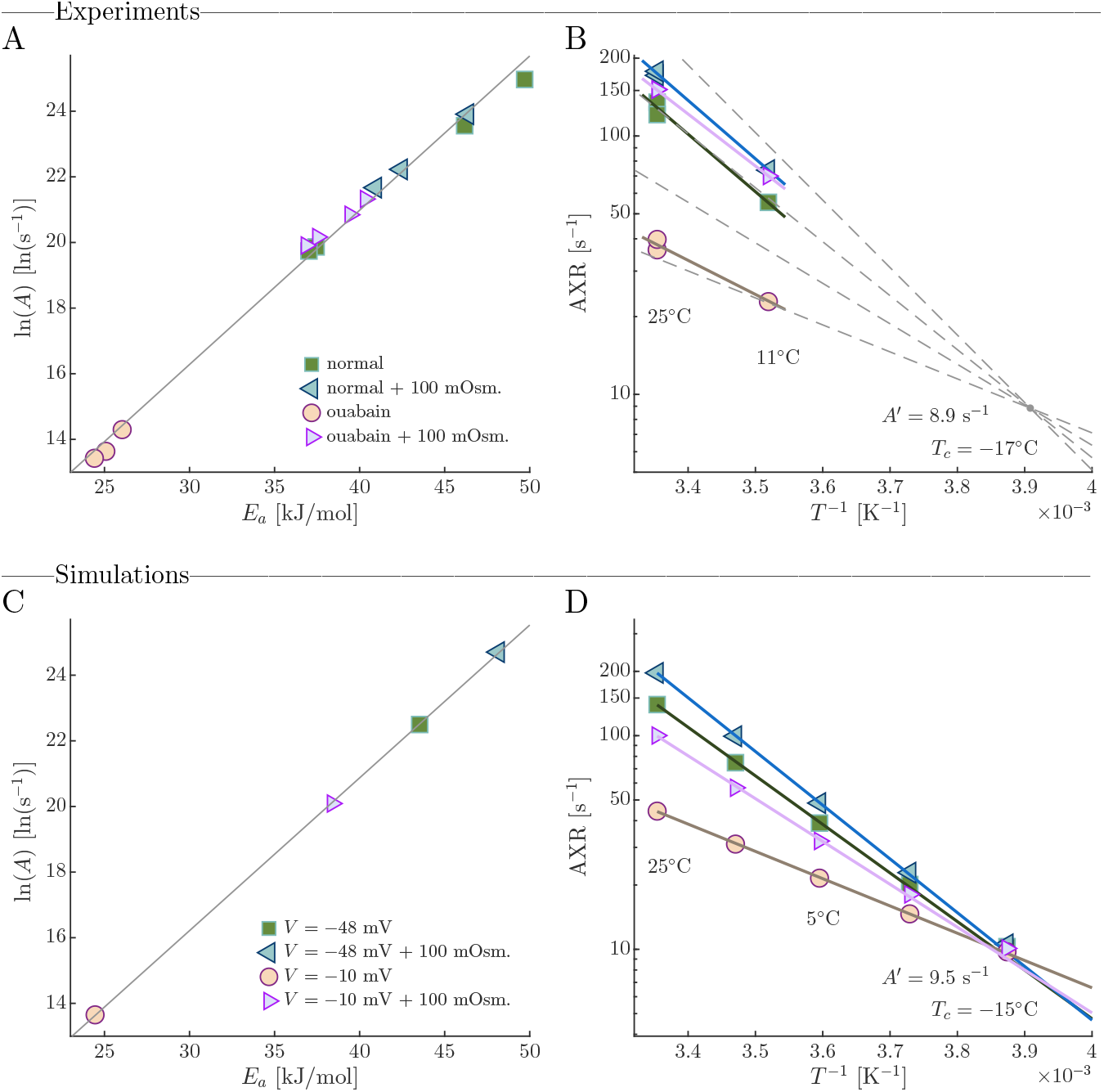
AXR increasing with activation energy follows enthalpy–entropy compensation. A) Experimental activation energies plotted vs. the natural log of pre-exponential factors for all samples show a linear dependence described by Eq. 2 with slope *m* = 0.470 ln (s^−1^)/(kJ/mol) (95% CI = 0.447, 0.493) and *y*-intercept ln (*A*′) = 2.18 ln (s^−1^) (95% CI =1.30, 3.06). This is a sign of EEC and suggests a common crossover temperature *T*_*c*_ = 1/(*m × R*) − 273.15 = −17.1 ^°^C (-29,-4.3). B) Semi-log plot of Eq. 3 for a range of *E*_*a*_ (dashed lines) along with the data from Fig. 5F. C) *E*_*a*_ vs. ln (*A*) estimated from 3XM simulations. D) 3XM predictions of AXR at temperatures between −25 and +25^°^C under voltage and osmolarity conditions modeling those studied experimentally (see legend in C). The modified Arrhenius model (Eq. 3) shows a common crossover temperature at *T*_*c*_ = −15^°^C (-20,-9.2).

While other explanations for EEC remain possible, we propose a multi-site exchange mechanism in which tonicity modulates AXR and the overall activation energy by shifting ECS and ICS volume fractions, thereby changing the relative prominence of exchange pathways with distinct kinetics and activation energies.(26) We test this hypothesis by simulating temperature-dependent DEXSY data using a three-site exchange model (3XM) with exchange between an ECS compartment and two ICS compartments (Fig. 8) (See Methods, SI, and Ref. 26). In the 3XM, transmembrane exchange between ECS and ICS is assigned a faster rate constant (*k*_*t*_) with a higher *E*_*a*_, whereas geometric exchange between ICS compartments (*k*_*g*_) is slower and has a lower *E*_*a*_. To demonstrate that the model can reproduce behavior similar to that observed experimentally, the *E*_*a*_ values were selected based on the extremes of the experimental observations, and the corresponding *A* values were chosen to match the fit in Fig. 7A. Thus, the parameterization is illustrative rather than independently derived. Tonicity alters ECS and ICS fractions via osmotic balance, which in turn changes DEXSY’s sensitivity to each pathway. Intuitively, sensitivity to any exchange pathway depends on the signal fractions of the exchanging compartments involved. Decreased ECS volume fraction reduces the sensitivity to transmembrane exchange, shifting sensitivity towards geometric exchange. As ECS volume fraction increases, sensitivity shifts toward the fast, high-*E*_*a*_ transmembrane pathway, causing both AXR and its apparent *E*_*a*_ to increase. The 3XM reproduces the observed EEC behavior, including similar values of *T*_*c*_ and *A*′ (Fig. 7C,D). Although *T*_*c*_ lies below 0^°^C and cannot be experimentally accessed, 3XM simulations show AXR converging at this point, where AXR = *k*_*t*_ = *k*_*g*_ = *A*′ (Fig. 7D).

**Fig. 8.**
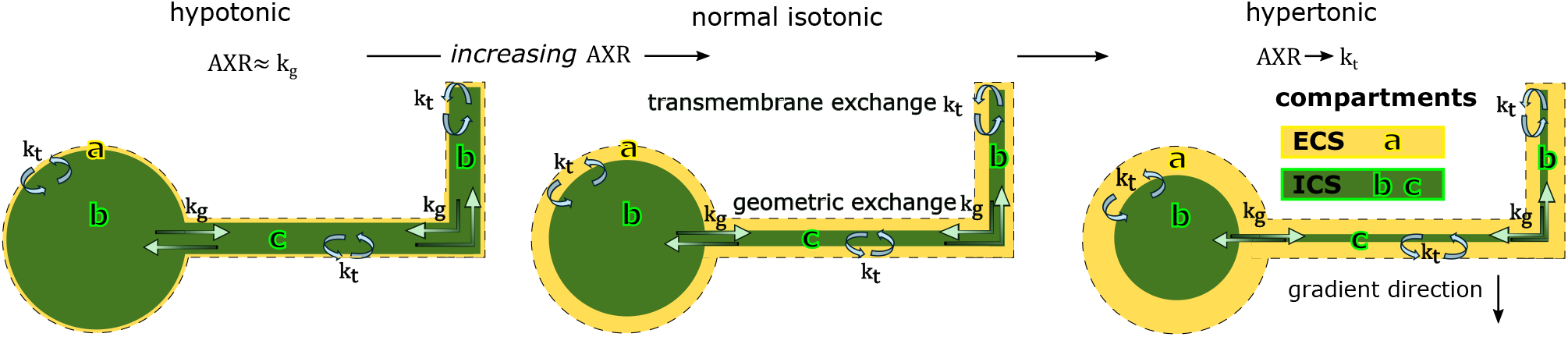
Depiction of the connection between tonicity and AXR in a three-site exchange model (3XM) for gray matter. Compartment **a** represents the ECS. Compartment **b** represents cell soma and processes oriented parallel to the gradient direction where water appears more mobile. Compartment **c** represents cellular processes oriented perpendicular to the gradient direction where water appears less mobile. Transmembrane exchange *k*_*t*_ occurs between the ECS and ICS compartments. Geometric exchange *k*_*g*_ occurs between ICS compartments. Tonicity modulates AXR by changing the ECS fraction and the relative proportion of water seen exchanging with ECS.(26)

EEC was able to describe AXR of all samples at all tonicities and temperatures, implying a model to estimate tonicity from AXR. In SI 3, we develop this model by following Psurek *et al*. (2008)(43) and calibrate it using data from the ouabain and ouabain + 100 mOsm conditions. For normal samples, the model predicts Δ*c* = 90 mOsm. This closely matches the +82 mOsm required to restore AXR in ouabain-treated samples determined by a linear fit of AXR versus sucrose concentration (Fig. S6). We interpret this as the change in tonicity associated with partitioned ions or, equivalently, intracellular impermeants and their countercations. Since no established method directly measures tonicity inside intact neural tissue, it is unfortunately not possible to validate this.

## Discussion

Na^+^/K^+^–ATPase inhibition by ouabain is considered a model for terminal depolarization of membrane potential(46, 47) which often occurs during traumatic brain injury and stroke.(48) During terminal depolarization, ions and water repartition, leading to hypotonicity, cell swelling, and dramatic ECS shrinkage.(49, 50) The abrupt decrease in both AXR and ADC_*y*_ (Fig. 1) is likely a consequence of this. Separating the effects of tonicity and voltage with sodium gluconate and ouabain showed the link between membrane potential and AXR is indirect, mediated by changes in tonicity (Fig. 2). The 3XM explains why AXR is more dramatically affected than ADC_*y*_.(26) These findings suggest AXR may serve as a sensitive biomarker for spreading depolarization, potentially outperforming ADC alone.(51)

Experiments decoupling tonicity from Na^+^/K^+^– ATPase activity showed that AWC is not significant in the *ex vivo* neonatal mouse spinal cord on 1–300 ms timescales. Inhibiting Na^+^/K^+^–ATPase activity did not reduce AXR when tonicity was maintained with impermeable osmolytes replacing semipermeable NaCl (Fig. 2). Osmolytes alone had little effect on AXR under normal conditions, despite likely reducing Na^+^/K^+^– ATPase activity,(39) but restored AXR under continued Na^+^/K^+^–ATPase inhibition (Figs. 3, 4). *E*_*a*_ recovered with AXR under conditions where AWC should not occur (Fig. 5).

While the 3XM could explain the link between water exchange and active ion transport reported in diffusion-based studies(6, 18, 20) it may not explain findings from studies using contrast agent-based methods(13, 14, 52–56) where geometric exchange is not expected. It also may not explain results on isolated cells(13, 52) where the ECS volume fraction is large and is unaffected by cell swelling or shrinking. There is a discrepancy between our results and theirs, which we currently cannot explain. For contrast agent-based experiments, multisite exchange could still arise from multiple exchanging compartments having different relaxation times. Changes in compartment volumes, even in the presence of contrast agents, could then result in the appearance that transmembrane exchange is affected. To resolve this issue, experiments controlling for the effect of tonicity while perturbing activity should be performed. Combined diffusion- and contrast agent-based studies separating the effects of activity and tonicity will also help understand this discrepancy.

Our data revealed a positive, tonicity-dependent correlation between AXR and *E*_*a*_, consistent with prior *ex vivo* studies comparing viable to fixed spinal cords(20) and viable to depolarized frog sciatic nerves.(53) The behavior was consistent with EEC. A model was developed based on EEC to estimate tonicity from AXR. While the model still needs further validation, the ability to non-invasively quantify tonicity in tissue is an important by-product of this work, since tonicity is a critical characteristic of the cellular state but is not measurable otherwise.

The 3XM was introduced as a minimal model to explain our results (Fig. 8).(26) Tonicity alters ECS volume fraction and hence the fraction of water molecules which DEXSY encodes as exchanging between ECS and ICS. This shifts the sensitivity of AXR between slower, low-*E*_*a*_ geometric exchange when the ECS fraction is smaller, and faster, high-*E*_*a*_ transmembrane exchange when the ECS fraction is larger. The sigmoidal curve in Fig. 4F traces this shift. The relative insensitivity of AXR at the lower and higher portions of this sigmoidal curve can explain the non-zero *y*-intercept in Fig. 1F insert and the small effect of 100 mOsm on normal samples in Fig. 3C,D, and suggests that below and above a certain ECS fraction the AXR is unaffected.

The slower exchange has *E*_*a*_ ≈ 25 kJ/mol, similar to water self-diffusion *E*_*a*_ ≈ 20 kJ/mol in pure water (40) and *E*_*a*_ ≈ 18 kJ/mol in aCSF. This is expected for geometric exchange involving hindered diffusion between intracellular compartments. Given the predominance of gray matter(28) and known microstructural features of the neonatal mouse spinal cord,(30, 31) such exchange could occur along branched cellular processes(16, 17) or between soma and processes.(25)

In contrast, the faster, but still passive exchange corresponds to a higher *E*_*a*_ ≈ 40 kJ/mol. This value is indicative of transmembrane water exchange, and not geometric exchange. Potential pathways for transmembrane exchange include channels, cotransporters, and the lipid bilayer. Water channels such as aquaporin would reduce *E*_*a*_ towards 20 kJ/mol(57) and hence play a minimal role. Ion cotransporters were also ruled out since it was shown by replacing NaCl with sucrose that it’s the tonicity and not the ions themselves that are important. The *E*_*a*_ value is consistent with passive diffusion across lipid bilayers.(21) With no other high *E*_*a*_ pathways existing in high abundance, we conclude the lipid bilayer to be the dominant transmembrane exchange pathway on 1 to 300 ms timescales in this system.

This mechanism is quantitatively plausible. Geometric exchange persists over broad and long timescales, both because it is mediated by diffusion which scales with 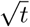 rather than exp(*t*) (16, 17) and because the microstroctural length scales are large. For instance, diffusion across typical branching processes lead to ≈ 80 to 0.1 s^−1^ for 5 to 145 *µ*m branch lengths(24)) and diffusion escape from soma to processes lead to ≈ 15 s^−1^ using Eq. 10 in Ref. 58 with an average 7 *µ*m radius soma connected to six 0.8 *µ*m radius processes(24). In comparison, transmembrane exchange rate constants are fast and predicted to exceed 100 s^−1^, based on SVRs of 4 to 16 *µ*m^−1^ in gray matter,(24) and permeabilities ranging from 70 to 400 *µ*m/s for polyunsaturated membranes and decreased by about 50% with 40% cholesterol content.(21, 22)

The 3XM predicts ADC and AXR to both depend on ECS volume fraction. We found ADC to correlate linearly with volume fraction metrics, but AXR showed a complex correlation (Figs. S2 and S5). Experiments show that ADC and AXR hold unique information during physiological vs. pathological cell swelling and shrinking (Fig. 4F). In essence ADC measures the global average of ADCs in every compartment, whereas AXR is related to volume fractions and eigenvalues of the exchange matrix.(26) Note also that compartments which do not exchange on the timescale of the measurement will affect ADC by their volume fraction but will not affect AXR.(59) While more possibilities exist, one explanation for this discrepancy between the 3XM and experiments is that the tissue is very heterogeneous, containing many more than three compartments, and the volumes of these compartments can change in a heterogeneous way. At this point, we believe that ADC and AXR are reporting different aspects of cell swelling.

These findings position DEXSY NMR as a powerful, noninvasive tool for probing steady-state water exchange and its biophysical and physiological underpinnings in viable tissue. More work is needed to understand the generality of these findings to more conventional gradient strengths and the *in vivo* human brain.(2–5) Nonetheless, future work is motivated by the possibility to infer tonicity and tissue state from combined measurements of AXR and ADC, to provide new biomarkers for monitoring normal function, development, aging, disease, and trauma without the need for contrast agents.

## Materials and Methods

All experiments were carried out in compliance with the *Eunice Kennedy Shriver* National Institute of Child Health and Human Development Animal Care and Use Committee under Animal Study Proposal (ASP) # 21-025. Neonatal mouse spinal cords were isolated from Swiss Webster wild type mice (Taconic Biosciences, Rensselaer, USA) between postnatal day 1 and 4 in a dissecting chamber perfused with low-calcium, high-magnesium artificial cerebrospinal fluid (aCSF), with the same composition as normal aCSF (listed below) but 0.5 mM CaCl_2_ · 2H_2_O and 6 mM MgSO_4_ · 7H_2_O, bubbled with 95% O_2_/5% CO_2_. After dissection, the spinal cord was placed in a RF coil in the sample chamber in normal calcium aCSF, composed of 128.35 NaCl, 4 KCl, 1.5 CaCl_2_ · 2H_2_O, 1 MgSO_4_ · 7H_2_O, 0.58 NaH_2_PO_4_ · H_2_O, 21 NaHCO_3_, 30 D-glucose (concentrations in mM), bubbled with 95% O_2_ and 5% CO_2_. aCSF flowed through the chamber at 7 mL/min. Stock solutions of the pharmacological Na^+^/K^+^–ATPase inhibitor ouabain (European Pharmacopeia Reference Standard) and osmolytes sucrose or mannitol (Sigma Aldrich) were prepared in media and added to the media reservoir to reach desired concentrations. Washing to media in which the 128.35 mM NaCl was replaced with 256.7 mM sucrose or 128.35 mM sodium gluconate (Sigma Aldrich) or back to normal aCSF media involved washing in an open loop with 100 mL of new media and then closing the loop. Sample temperature was monitored by a fiber optic sensor (PicoM, Opsens Solutions Inc., Québec, Canada) and controlled by a chiller that circulated water through the chamber base. Samples were normally maintained at 25 ± 0.2^°^C. Temperature perturbations involved switching to a second chiller which maintained the sample at 11 ± 0.2^°^C (5 min equilibration). See Ref. 20 for more details.

NMR experiments were performed with a single-sided permanent magnet(60) (PM10 NMR MOUSE, Magritek, Aachen, Germany) at field strength B_0_ = 0.3239 T and gradient strength *g* = 15.3 T/m. NMR experimental protocols involved looping through 11-minute sets of diffusion experiments, rapid diffusion exchange experiments, and *T*_1_ saturation recovery experiments. Diffusion experiments were performed using a standard spin echo sequence(61) and 4-step phase cycle.(62) The encoding time, *τ* of the spin echo was varied linearly from 0.05 to 3.3 ms over 22 steps with 4 scans per *τ* . This corresponds to *b*-values ranging from 0.001 to 400 ms*/µ*m^2^ where *b* = 2/3*γ*^2^*g*^2^*τ* ^3^ and *γ* is the gyromagnetic ratio. Points two through four (*τ* = 0.205 to 0.514 ms, *b* = 0.096 to 1.5 ms*/µ*m^2^) were fit with *I*(*b*) = *I*_0_ exp ( − *b* ADC) to estimate ADC. Rapid diffusion exchange experiments were performed following the Diffusion Exchange Ratio (DEXR) method (63) using a DEXSY sequence involving two spin echoes separated by a mixing time *t*_*m*_ in the presence of a static magnetic field gradient.(32) A 4-step phase cycle selects only the desired DEXSY coherence transfer pathway.(64) Experiments were performed with (*τ*_1_, *τ*_2_) combinations (0.200, 0.735) ms and (0.593, 0.580) ms, 8 scans per combination, and *t*_*m*_ values [0.2, 1, 2, 4, 7, 10, 20, 40, 80, 160, 300] ms. The (0.200, 0.735) signals were fit with a biexponential model. The (0.593, 0.580) signals were divided by the biexponential model and then fit with *I*(*t*_*m*_) = *β*_1_ exp (− *t*_*m*_AXR) + *β*_3_ to estimate the apparent exchange rate constant, AXR (similar to Method 3 in Ref. 65). See Refs. 32 and 20 for more details.

Paired-sample t-tests, two-sample t-tests, and one-way ANOVA were performed to test for effects, differences between means of treatment groups, and recovery, respectively, with *p* < 0.05 considered significant.

In the 3XM, volume fractions were defined based on an osmolarity balance and fixed total volume (Eq. S8). The osmolarity balance includes trapped impermeants, small osmolytes, and ions partitioned by the voltage. We used −48 mV to model normal conditions (based on intracellular recordings (66)) and −10 mV to model Na^+^/K^+^–ATPase inhibition by ouabain (based on recordings under terminal depolarization (49)). The initial ECS fraction was set to 0.3, based on realtime iontophoresis measurements in a neonatal mouse spinal cord slice model (23). AXR was modeled by simulating DEXSY data using an operator formalism(67) and then fitting the DEXR model described above. See SI 4 and Ref. 26 for more details.

## Data, Materials, and Software Availability

All data and code used to generate figures has been made publicly available at https://data.mendeley.com/datasets/rv3jmhb7wz

## ACKNOWLEDGMENTS

This work was partially funded by the Department of War in the Military Traumatic Brain Injury Initiative (MTBI^2^) under award HU0001-24-2-0051. NHW, RR, TXC, and PJB were also supported by the IRP of the NICHD, NIH (Project numbers (PN): 1ZIAHD008971-07; 1ZIAHD008972-07). JAR was supported by an NIGMS Postdoctoral Research Associate Training (PRAT) Program Fellowship Award (PN: 1FI2GM150429-01; 1ZIEGM000002-18). Chat GPT-5.3 was used to help edit for grammar and clarity. We thank MJ O’Donovan and M Falgairolle for helpful discussions regarding the neonatal mouse spinal cord model, CS Springer, T Barbara, and V Kiselev for carefully reviewing the work, RW Pastor, AM Berezhkovskii, and SM Bezrukov for input on lipid membrane and channel permeability, and JF Douglas for bringing EEC to our attention.

## Supporting Information

### Supporting Information Text

#### 1. ADC provides a proxy for diffusion and DEXSY-derived volume fractions

Simultaneous realtime measurements were used to test for correlations between the metrics ADC_*y*_, AXR, *f* (restricted water fraction), *c*, and *f*_*ng*_ (non-Gaussian signal fraction). ADC_*y*_, *f*, and *c* arise from static gradient spin echo (SGSE) diffusion experiments. ADC_*y*_ measures the average mobility of all water molecules in the sample on the timescales of diffusion encoding (here between 0.20 and 0.51 ms). *f* and *c* are measured from fitting *I/I*_0_ = *f* exp (− *τc*) to the SGSE diffusion signal attenuation at *b* ≥ 60 ms*/µ*m^2^ and are associated with localization or motional averaging of water near surfaces on the length scale of the gradient dephasing length *ℓ*_*g*_ (see Williamson, et al., Magn. Reson. Lett., 2023) Here, with the extremely strong *g* = 15.3 T/m static gradient produced by the low-field single-sided magnet, *ℓ*_*g*_ = 0.8 *µ*m (at 25^°^C). *c* is the rate constant for the decay of this localized or motionally-averaged component as a function of echo time. *c* is related to a restriction length scale in the case of motional averaging, or *D*_0_ in the case of localization, and also affected by exchange. ADC is a standard measurement used clinically (although with longer diffusion timescales between 20 and 100 ms), whereas *f* and *c* require gradient strengths not currently accessible with the pulsed gradients of clinical systems. *f*_*ng*_ is derived from the DEXSY data (see Eq. 19 in Cai, et al., J. Magn. Reson., 2024). Correlations between ADC and *f* have not been studied before, although they are often inferred in two-component models for ADC. Such models may include a hindered component and a restricted component with apparent diffusion coefficients *D*_1_ and *D*_2_ respectively, leading to ADC = (1 − *f* )*D*_1_ + *f D*_2_. Hindrances reduce *D*_1_ below the diffusion coefficient of freely diffusing water, *D*_0_ = 2.15 *µ*m^2^/ms at 25^°^C. Water that is restricted on length scales similar to or smaller than *l*_*g*_ will have *D*_2_ ≪ *D*_1_, leading to ADC = (1 − *f* )*D*_1_ in the limit that *D*_2_ = 0. Note that the slope and intercept here are equal to *D*_1_, the diffusion coefficient of the more mobile compartment. We use correlations to test this model and understand what the metrics are measuring in this system.

We find strong linear correlations between ADC_*y*_ and *f* (Figs. S2D and Figs. S5D). The correlation is the same regardless of the type of perturbation (with/without ouabain and osmolytes). Slopes and *y*-intercepts are similar to *D*_0_ consistent with the model. These results are consistent with the picture that *f* is associated with effectively immobile water and ADC_*y*_ is a proxy for *f* . This suggests that both ADC_*y*_ and *f* report cellular swelling and shrinking.

ADC_*y*_ and *f*_*ng*_ are strongly correlated under normal conditions, with slope and intercept slightly above *D*_0_, but not correlated after ouabain treatment (Figs. S2F). During osmolyte perturbations, the correlation appears strongest when ΔADC_*y*_ and Δ*f*_*ng*_ are compared (Figs. S5F). The slopes are very close to 1, as predicted by the simple model. Deviations here are expected, since ADC_*y*_ and *f* are global measures averaged over the entire sample, whereas *f*_*ng*_ is a local measure averaged over exchanging compartments.

#### 2. Discussion about differences observed during the wash

Osmotic perturbations with 100 mM sucrose and with variable concentrations up to 200 mM yielded consistent results overall (Fig. 4C,E). Differences were observed during the wash from hypertonic media to normal media on samples not treated with ouabain. AXR recovered to stable values slightly below baseline after washing out the +100 mOsm solution, but dropped to low levels, similar to ouabain-treated values, after washing out the +200 mOsm solution. (compare Fig. 3C to Fig. 4A). It appears that the samples maintained homeostasis during the wash in the first case but not in the second. This could be because of the larger osmotic change in combination with a longer experiment time. 200 mM is more than cell RVI mechanisms can physiologically balance given the limited NaCl and KCl in the aCSF and may have damaged the cells. Additionally, intracellular penetration of sucrose may have caused rebound swelling when washed to normal media. For reference, the wash occurred at roughly 120 minutes in the 100 mM case and at roughly 300 minutes in the 200 mM case.

#### 3. Derivation of EEC-based model to predict tonicity

In this section we derive a model relating a rate constant *k* (such as AXR) to tonicity. The model is based on an application of transition state theory to the dynamics of mixtures. We first define the the enhancement of exchange with tonicity relative to a state of minimum (hypo)tonicity as *θ* = *k*_*t*_*/k*_0_. Within transition state theory, *θ* = exp (− *δG*_*t*_*/RT* ) where *δG*_*t*_ = |*G*_*t*_ − *G*_0_| is the change in Gibbs free energy associated with the difference in osmolarity of impermeable solutes in the extracellular space which increase the tonicity (hence the subscript t). Changes in Gibbs free energy can be decomposed into enthalpic and entropic contributions by its thermodynamic definition *δG*_*t*_ = *δE*_*at*_ − *TδS*_*t*_. Then

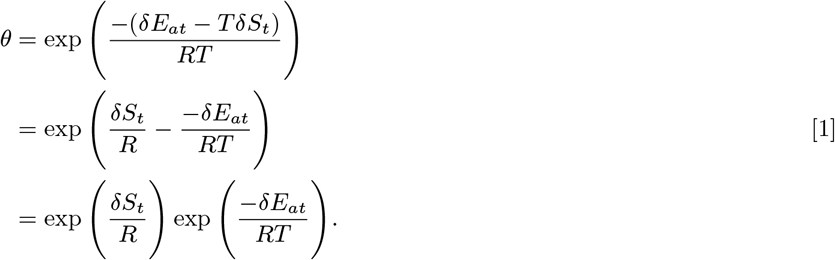

To become the Arrhenius model, we explicitly define the connections between the change in entropy and the preexponential factors exp (*δS*_*t*_*/R*) = *A*_*t*_*/A*_0_, and between the change in enthalpy and the activation energies *δE*_*at*_ = *E*_*at*_ − *E*_*a*0_. Here, *A*_*t*_ and *E*_*at*_ are tonicity-dependent and *A*_0_ and *E*_*a*0_ are evaluated when the system is at a a state of minimum (hypo)tonicity, defining *c* = 0. Then

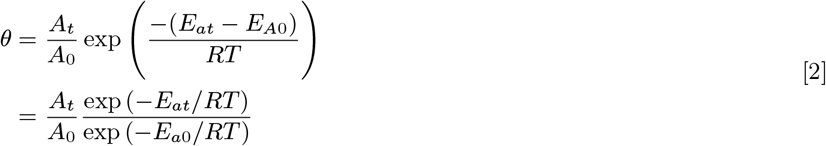

Now the effect of tonicity on exchange can be considered in terms of its separate effect on *δE*_*at*_ and *δS*_*t*_.

The analytical functions for *δE*_*at*_ and *δS*_*t*_ with respect to the osmotic concentration can be obtained by Taylor expansions,

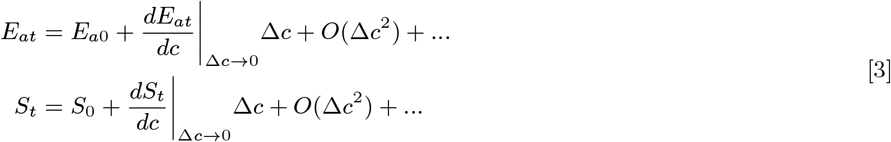

where *O*(Δ*c*^2^) + … refer to second- and higher-order terms and |_Δ*c* → 0_ refers to the term being evaluated in the limit of Δ*c* → 0, i.e., *c*_*t*_ → *c*_0_. We limit ourselves to a first-order approximation so that a two-parameter model can be obtained, but with the downside that the model loses validity as Δ*c* increases above dilute concentrations. We also drop the evaluated symbol for brevity (although still implied). Then

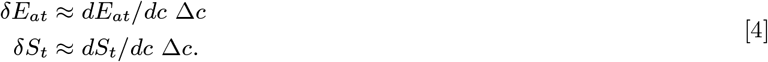

These two parameters can be calculated from measurements of *E*_*a*_ and *A* at *c* = *c*_0_ and *c*_1_ with Δ*c* = *c*_1_ by using the relations 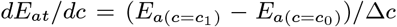 and 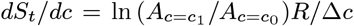. These are related to virial coefficients for the entropy and enthalpy of activation by [*E*_*at*_] = (*dE*_*at*_*/dc*)*/R* and [*S*_*t*_] = (*dS*_*t*_*/dc*)*/R*. These definitions are combined with Eq. 1 to obtain the final relationship

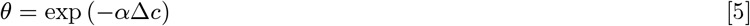

where

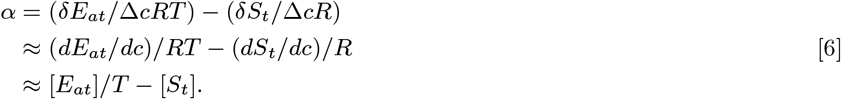

From this model, it is also possible to calculate the tonicity Δ*c* as

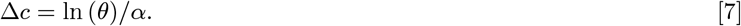

For our system, by comparing *E*_*a*_ and *A* of samples after treatment with ouabain to samples after treatment with ouabain and 100 mM sucrose (*c* = *c*_0_ and *c* = *c*_1_, respectively), we find [*E*_*at*_] = 16.1 K/mM and [*S*_*t*_] = 0.0686 1/mM, leading to *α* ≈ − 0.0146 1/mM at 25^°^C. Based on *k*_*t*_ under normal and *k*_0_ from ouabain-treated conditions *θ* = 130/35, and Eq. 7 predicts Δ*c* = 90 mOsm.

#### 4. Additional PLM and 3XM details

The pump–leak model (PLM) uses the finite difference method to approximate the fluxes of ions based on their governing differential equations for mass conservation. Na^+^ and K^+^ transport actively with a 3:2 stoichiometry. Na^+^, K^+^, and Cl^−^ transport passively based on their electrochemical potential. The net charge of the intracellular impermeants is *z* = − 1. Water permeability is assumed to be much higher than ion permeability and is not modeled directly. Instead, the cell changes volume so that the intracellular osmolarity matches the extracellular osmolarity at the end of each timestep. Voltage is modeled based on the net charge of the intracellular ions and the (constant) capacitance of the membrane. Unless stated otherwise, simulation parameters were the same as in Kay (2017). Volumes and voltages were taken as the values obtained at the final timestep. This time was sufficiently long for systems to reach steady-state (if there existed a stable steady-state), as determined by volume and voltage not changing when total time was increased by a factor of 10.

An analytical equation for the steady-state ECS fraction (*f*_*o*_) was developed based on an osmotic balance which includes the contribution of an uncharged, trapped ECS osmolyte (*x*_*o*_);

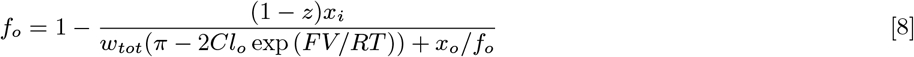

where *x*_*i*_ is number of moles of intracellular impermeants (i.e., solutes which are trapped in the ECS or ICS that can not cross the plasma membrane or communicate with the bathing media), *π* is the concentration of small osmolytes in the ECS (i.e., solutes that can not cross the plasma membrane but can communicate with the bathing media), *Cl*_*o*_ is the concentration of extracellular chloride, *F* is the Faraday constant, *R* is the ideal gas constant, and *T* is the absolute temperature. *π* and *Cl*_*o*_ are assumed equal to that of the bathing media. The ECS and ICS volumes sum to a fixed total volume, *w*_*tot*_. While this model does not explicitly account for non-osmotic (e.g. hydrostatic and mechanical) pressure gradients which could become large when cells swell and push against one-another or when cells shrink and push on intracellular organelles, the trapped ECS and ICS impermeant parameters act as placeholders to effectively model this buildup of pressure in cases of extreme swelling or shrinking. This equation does not model ion flux and instead assumes a value for the voltage (*V* ). We used −48 mV for normal conditions and −10 mV for the conditions modeling Na^+^/K^+^–ATPase inhibition with ouabain. First, the PLM was used to find the steady-state cell volume predicted under normal conditions. This, along with *f*_*o*_ = 0.3 defined *w*_*tot*_. Then *x*_*o*_ was arbitrarily set to 1/50th of the moles of intracellular impermeants *x*_*i*_. Then Eq. 8 was used to predict ECS fraction *f*_*o*_ and osmolarities of intracellular and extracellular impermeants at varying osmolarities for the *V* = −48 mV and −10 mV conditions. A three-site exchange model (3XM) associated with an ECS compartment (**a**) and two ICS compartments (**b** and **c**) was developed for predicting ADC and AXR. Eq. 8 was used to define the fraction of compartment **a** (*f*_*c*_ = *f*_*o*_). For simplicity, the fractions of the two ICS compartments were set equal, *f*_*b*_ = *f*_*c*_ = (1 − *f*_*a*_)/2. The ECS compartment and a more mobile ICS compartment were both given ADC_*a*_ = ADC_*b*_ = 1 *µ*m^2^/ms. A less mobile ICS compartment was given ADC_*c*_ = 0.1 ms*/µ*m^2^. Varying ADC values of the compartments affected the overall ADC, the size of effects on ADC, and the correlation between ADC and AXR but did not affect the overall interpretations or the EEC relationship. ADC was modeled as the sum of the ADC values of each compartment multiplied by their volume fraction, ADC = *f*_*a*_ADC_*a*_ + *f*_*b*_ADC_*b*_ + *f*_*c*_ADC_*c*_. AXR was modeled by simulating data from the DEXSY experiment and then fitting the exchange model to the data. DEXSY signals were modeled using an operator formalism **S** = [1, 1, 1] ∗ *OD*_2_ ∗ *OE* ∗ *OD*_1_ ∗ **S**_0_, ignoring relaxation, where *OD*_2_, *OE*, and *OD*_1_ are the operators for the second diffusion encoding, mixing time, and first diffusion encoding of the DEXSY pulse sequence, respectively. This involved multiplication of matrix exponentials which used the function expm() in MATLAB 2024a. The operators for the first and second diffusion encoding blocks were *OD*_1,2_ = expm(−*b*_1,2_**D**) with **D** = [ADC_*a*_, ADC_*b*_, ADC_*c*_]′. ∗ eye(3), where eye(3) is a 3 × 3 identity matrix. Equilibrium magnetization was **S**_0_ = [*f*_*c*_, *f*_*b*_, *f*_*a*_]′. The exchange operator was *OE* = expm(− *t*_*m*_**K**) with exchange matrix

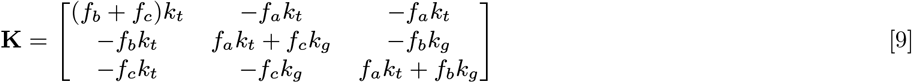

including transmembrane exchange (*k*_*t*_) and geometric exchange (*k*_*g*_ ) pathways. *k*_*t*_ and *k*_*g*_ values were determined by the Arrhenius model (Eq. 1) with *T* = 298.15 K or varied in the simulation, using *E*_*a*_ = 52 kJ/mol and ln (*A*) = 26.62 s^−1^ for *k*_*t*_ and *E*_*a*_ = 22 kJ/mol and ln (*A*) = 12.52 s^−1^ for *k*_*g*_ . These ln (*A*) values were defined for each *E*_*a*_ using Eq. 2 and the slope and intercept values presented in the caption of Fig. 7. Parameters (timings, gradient, *b*_1_ and *b*_2_ protocol, mixing times, etc.) were set based on values used experimentally. AXR values were estimated using the same DEXR model analysis method described in the “NMR hardware, experimental protocol, and analysis methods” section.

**Fig. S1.**
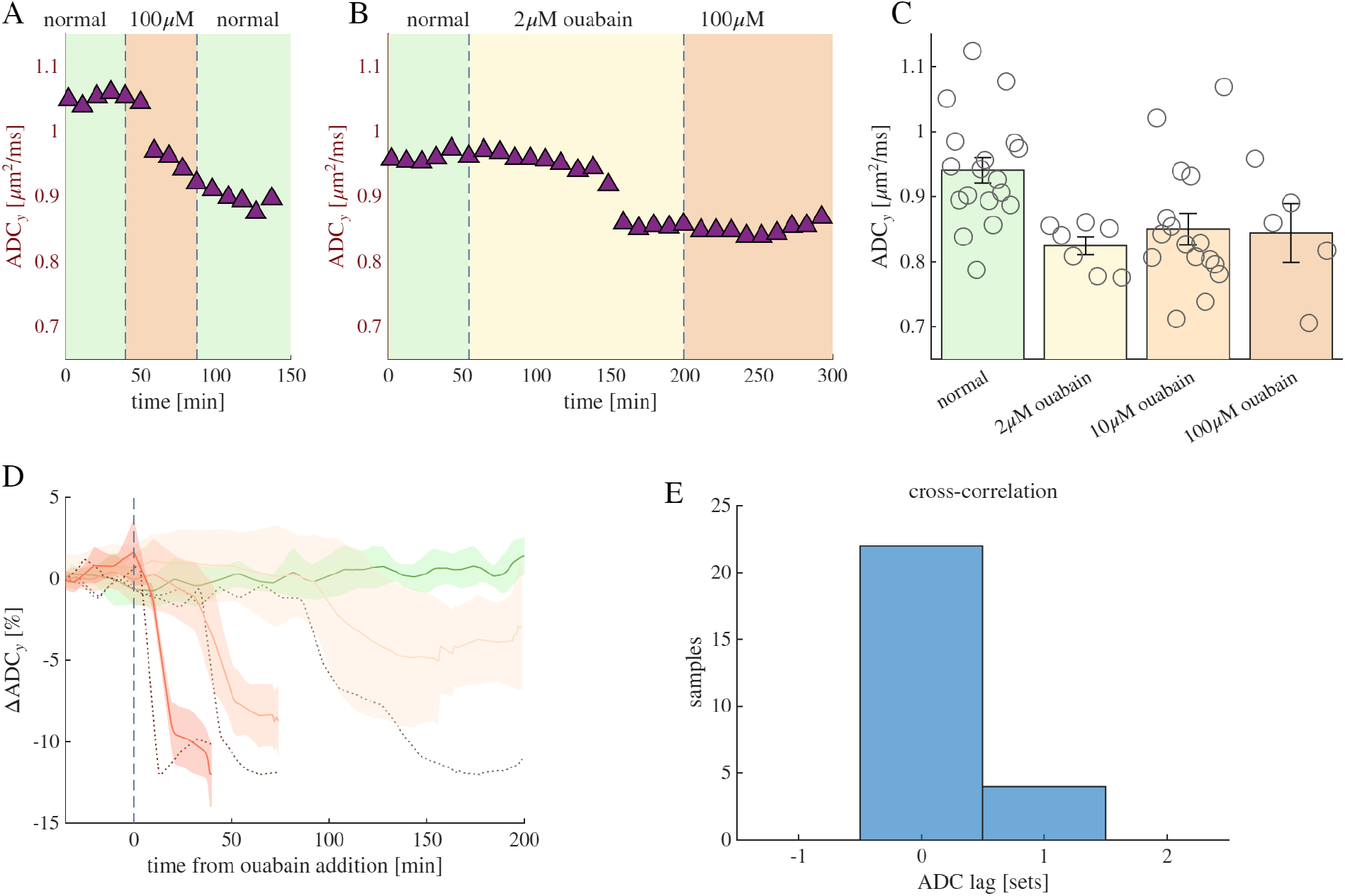
ADC_*y*_ and AXR are affected by ouabain simultaneously. A–C) data reproduced from Fig. 1, but presenting the absolute values of ADC. Note the additional scatter between samples in (C) compared to when normalized by baseline values in Fig. 1C. D) Mean (solid lines) and standard deviation (shaded regions) of realtime ADC_*y*_ measurements from a control group under normal conditions (green) and groups treated with 2, 10, and 100 *µ*M ouabain (lightest to darkest colors) plotted against the time after addition of ouabain. Dotted lines show the mean of AXR for each group, scaled to the maximum and minimum of the mean of ADC_*y*_ for comparison (refer to Fig. 1 for values) E) Histogram of the shift between the drop in ADC_*y*_ relative to the drop in AXR from a cross correlation analysis of 26 out of 28 samples for which there was a significant correlation between ADC_*y*_ and AXR (*cc >* 0.2, *p* < 0.05. Each set took 11 minutes to acquire and ADC_*y*_ measurements preceded AXR measurements. Analysis shows that ADC_*y*_ dropped either during the same set or the following set after AXR dropped, showing that ADC_*y*_ and AXR both dropped within 11 minutes of each other.

**Fig. S2.**
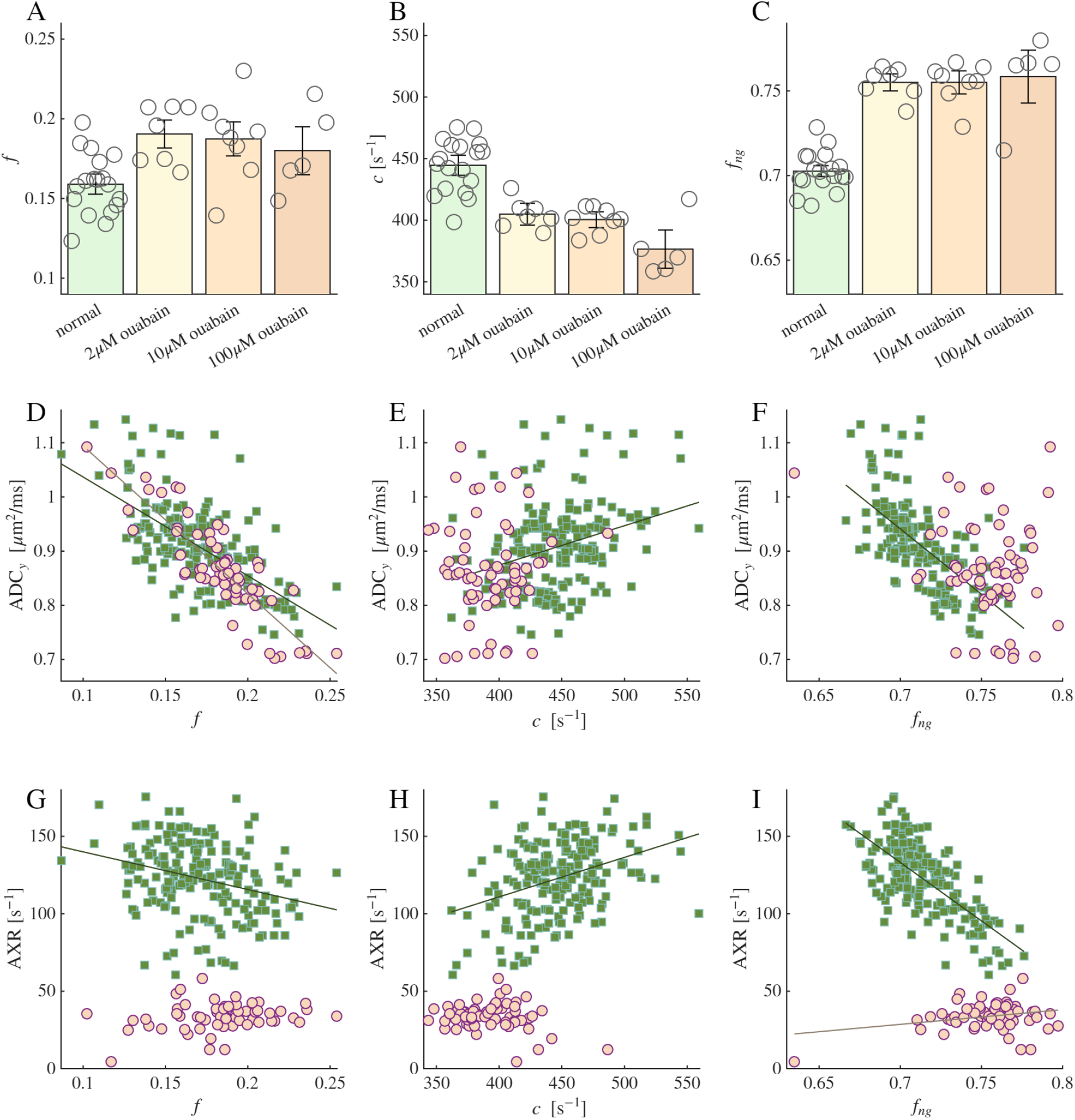
Additional metrics and correlations from the effect of ouabain treatment. A-C) Effect of ouabain on restricted fraction *f* and *c* from the intercept and slope of the SGSE diffusion signal at long times, and on the non-Gaussian signal fraction *f*_*ng*_ from the DEXSY data. D-I) Correlation of *f, c*, and *f*_*ng*_ vs. ADC_*y*_ and AXR. Data was grouped based on whether measurements occurred before (green squares) or after (pink circles) ouabain affected the exchange rate and dropped it below 60 *s*^−1^. Linear fits are plotted when correlation is significant (*p* < 0.05). In (D), the data is significantly correlated for both groups (*p* < 0.001) with Pearson correlation coefficients (cc=) -0.64 before ouabain effect and cc=-0.87 after ouabain effect. Linear fits yielded ADC_*y*_ = −1.82*f* + 1.22 *µ*m^2^/ms before ouabain effect and ADC_*y*_ = −2.72*f* + 1.36*µ*m^2^/ms after ouabain effect. In (F), the correlation is significant before the effect of ouabain, with (*p* < 0.001), cc=-0.65 and ADC_*y*_ = −2.4*f*_*ng*_ + 2.6 *µ*m^2^/ms, but is not significant after the effect.

**Fig. S3.**
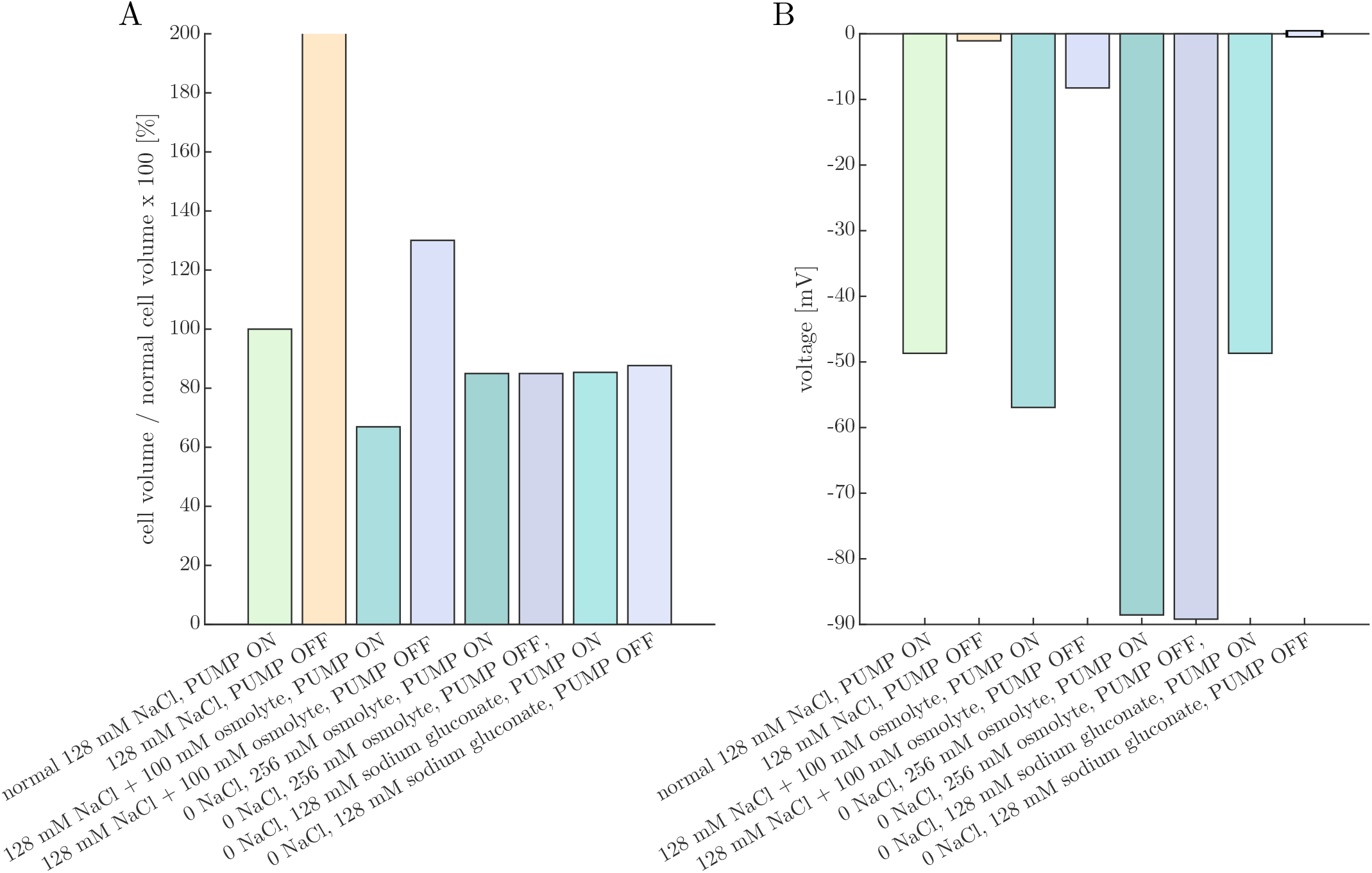
Pump-leak model (PLM) provides predictions of cell volume and voltage under conditions studied experimentally. A) Percent change in cell volume and B) voltage predicted under various Na^+^/K^+^–ATPase PUMP ON or OFF and media conditions after the PLM has run to steady-state (if applicable). With the 128 mM NaCl (and 4 mM KCl) media and PUMP OFF condition, the cell continues to swell and does not reach a steady-state. Addition of an osmolyte (+100 mOsm) reduces the volume and hyperpolarizes the cell in the PUMP ON condition, and stabilizes the volume but does not recover the voltage with the PUMP OFF. The cell volume is reduced to a similar level in both the PUMP ON and PUMP OFF conditions when NaCl is replaced by either sucrose or sodium gluconate (modeled as uncharged and monovalent anionic impermeants in the bathing media, respectively). However, while voltage is similar between the PUMP ON and PUMP OFF conditions when NaCl is replaced by sucrose, full depolarization to V = 0 is predicted in the PUMP OFF condition with NaCl replaced by sodium gluconate.

**Fig. S4.**
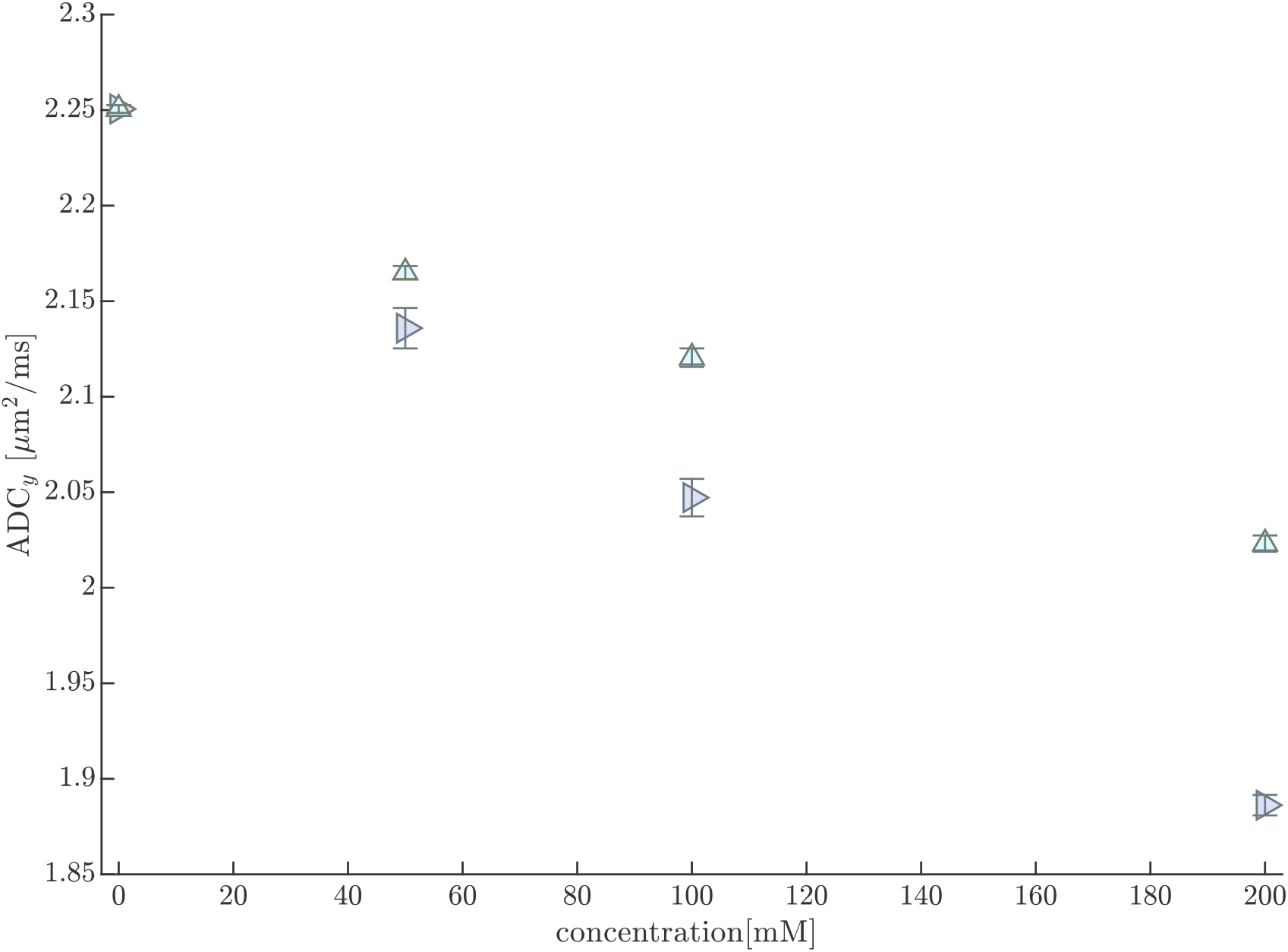
Concentration-dependent effects of sucrose and mannitol on ADC of pure aCSF. Spin echo diffusion measurements were performed on aCSF at varying concentrations of sucrose (pink rightward-pointing triangles) or mannitol (aqua upward-pointing triangles) at 25^°^C without media circulation. At least three measurements were performed at each concentration using 8 *τ* values spaced linearly from *τ* = 105 to 435*µ*s (*b* = 0.0129 to 0.9159ms*/µ*m^2^), TR=2 s, and 2000 echoes. all other parameters were the same as standard diffusion measurements performed on spinal cords. Sucrose affects diffusion of water more than mannitol because it is a larger molecule (increased obstruction) and affects viscosity more (increased hydrodynamic interactions). This leads to the conclusion that the compounds’ osmotic effects on tissue (cell shrinkage increasing ADC_*y*_ ) are more significant than their viscous effects (reducing ECS diffusivity). Their viscous effects may become apparent at higher concentrations, e.g. at 200 mM, where the average ADC of samples treated with sucrose is less than those treated with mannitol, although not significantly.

**Fig. S5.**
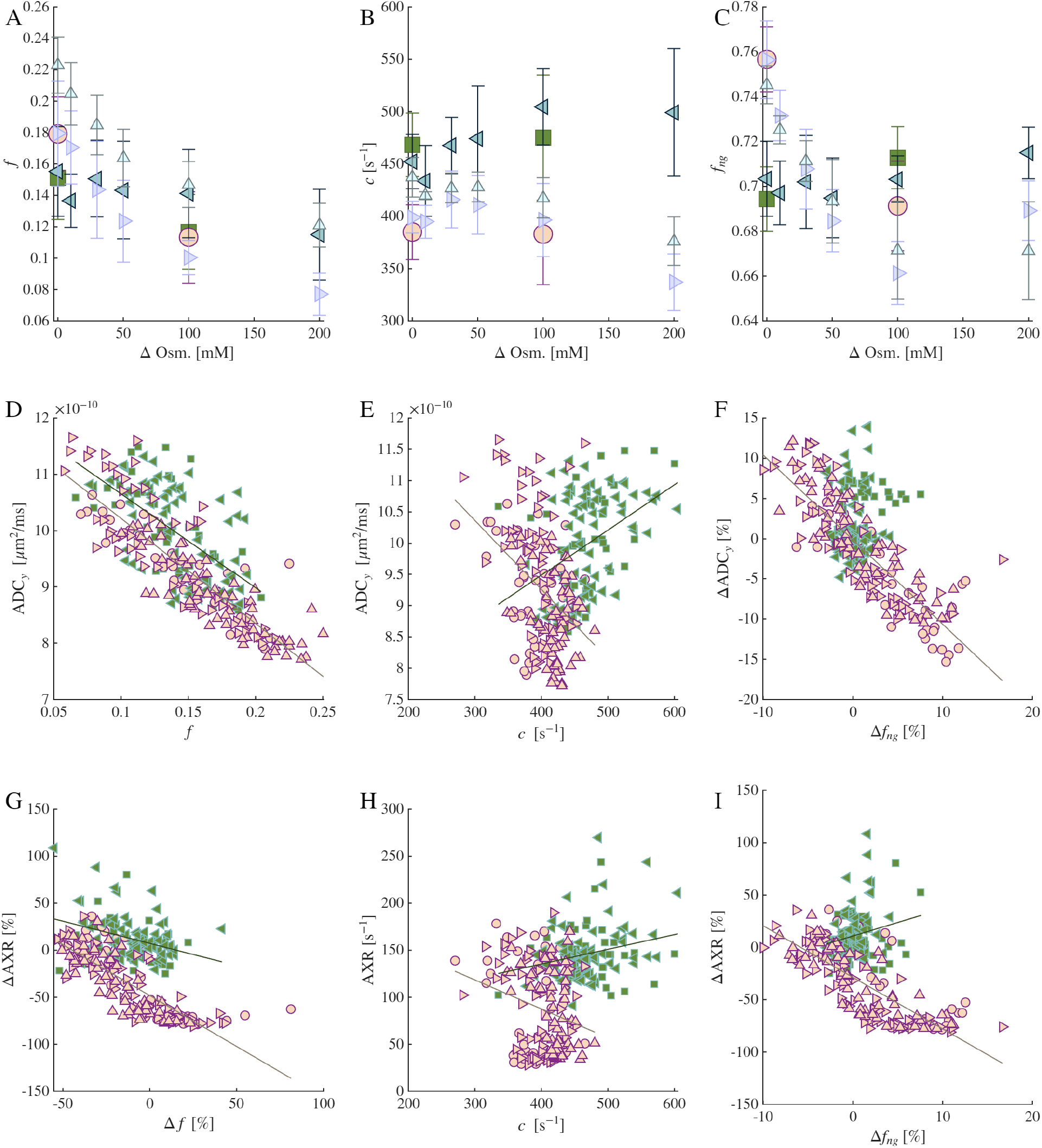
Additional metrics and correlations from the effects of osmolytes without and with ouabain treatment. A-C) Effect of osmolytes on *f* and *c* from the intercept and slope of the SGSE diffusion signal at long times, and on the non-Gaussian signal fraction *f*_*ng*_ from the DEXSY data. Refer to Fig. 1 for legend. D-I) Correlation of *f, c*, and *f*_*ng*_ vs. ADC_*y*_ and AXR. Data was normalized in cases where it tightened the correlation. In (D), cc= -0.61 without ouabain and cc=-0.87 with ouabain. Linear fits yielded ADC_*y*_ = − 1.67*f* + 1.23 *µ*m^2^/ms without ouabain and ADC_*y*_ = − 1.87*f* + 1.21*µ*m^2^/ms with ouabain (note the slope slightly reduced from *D*_0_). in (F), the correlation was insignificant without ouabain. With oabain, the correlation was significant (*p* < 0.001) with cc=-0.85 and linear fits yielded ΔADC_*y*_ = − 1.05Δ*f*_*ng*_ + 0.11 % (note the slope near 1). Note the sigmoidal correlations between Δ*f* vs. ΔAXR and Δ*f*_*ng*_ vs. ΔAXR

**Fig. S6.**
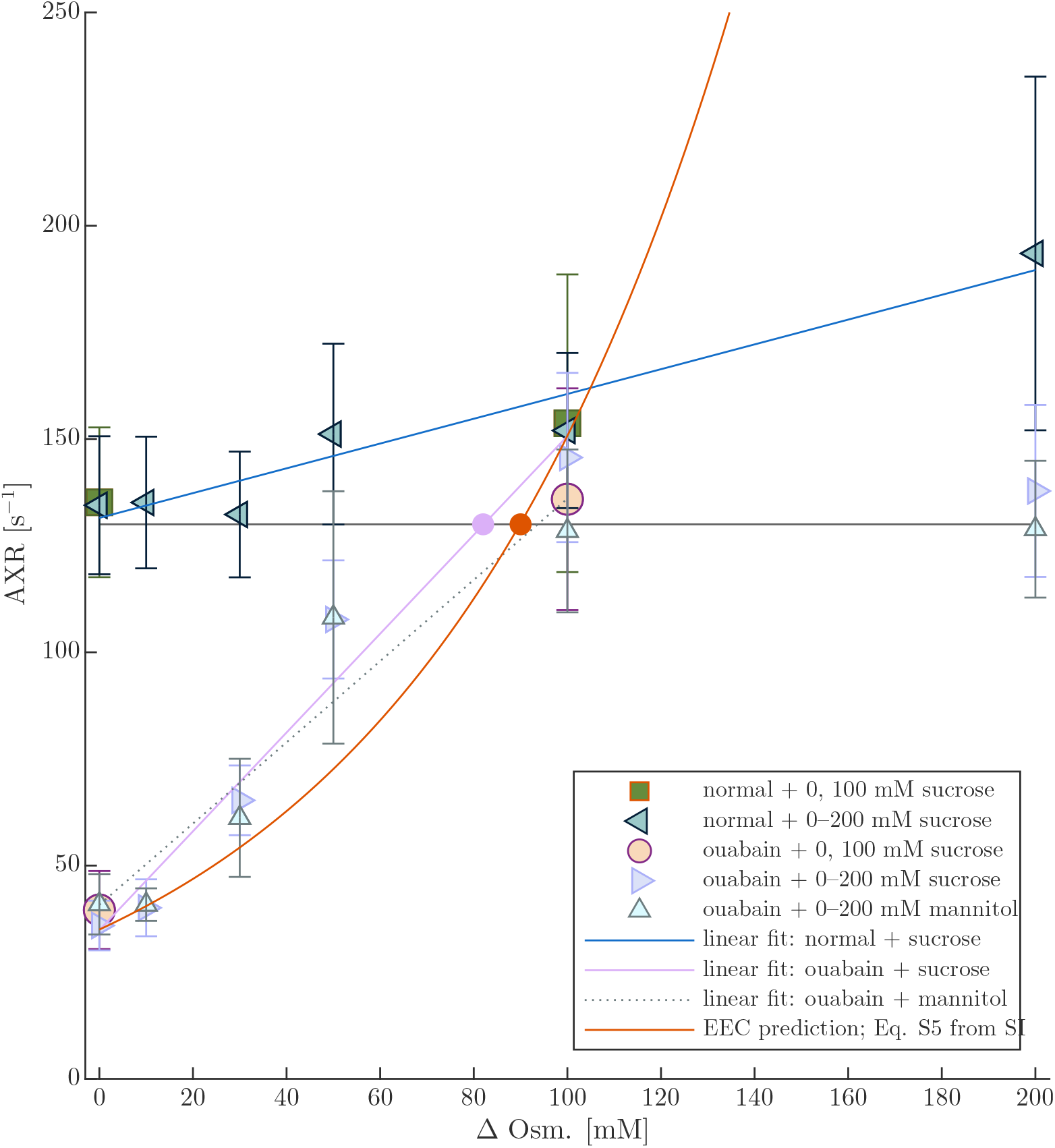
Linear fits and EEC prediction for osmotic dependence of AXR. Data reproduced from Fig. 4E from the main manuscript, with linear fits of the entire range (for normal + sucrose) or of the range from 0 to 100 mM for ouabain + sucrose and ouabain + mannitol, and the EEC prediction from Eq. 5 in SI section 3. The fit of the 0 to 100 mM for ouabain + sucrose data and the EEC prediction predict the AXR to equal the normal value (130 s^−1^) at 82 and 90 mOsm., respectively, shown by the filled circles.

